# Studies of all-*trans* retinoic acid transport in myopigenesis

**DOI:** 10.1101/2025.01.04.631331

**Authors:** Saptarshi Chatterjee, Ankana Roy, Jianshi Yu, A. Thomas Read, Melissa R Bentley-Ford, Machelle T. Pardue, Maureen A. Kane, M. G. Finn, C. Ross Ethier

## Abstract

**Purpose:** Myopia incidence is increasing globally. All-*trans* retinoic acid (atRA) is important in myopigenic retinoscleral signaling, motivating research on its ocular transport. However, atRA’s weak autofluorescence limits its direct visualization in tissues. Further, atRA is hydrophobic and must bind to protein carriers for transport. We assessed a fluorescent analog of atRA (LightOx^TM^14, CAS:198696-03-6, referred as ‘floRA’), as an experimentally accessible atRA surrogate by: (i) evaluating its binding to carrier proteins and (ii) visualizing its distribution in ocular tissues.

**Methods:** *Binding:* We assessed atRA-carrier protein binding using fluorescence quenching assays with bovine serum albumin (BSA), high density lipoprotein (HDL), apolipoprotein A-I (Apo A-I) and retinol binding protein 4 (RBP4).

*Direct visualization:* Wild-type C57BL/6J mice were euthanized, eyes enucleated, and wedges containing sclera and choroid incubated for specific durations in 50 μM floRA+BSA. The wedge centers were cryo-sectioned and counterstained for nuclei. Fluorescent micrographs were acquired and analyzed using ImageJ.

**Results:** Association constants (K_a_) for atRA and floRA binding to carrier proteins were similar and ranged from 2-13 × 10^5^ M^-1^, indicating non-specific binding. floRA could be visualized in sclera and choroid, yet showed significant spatial heterogeneity (enhanced fluorescence often colocalizing with nuclei).

**Conclusions:** floRA is a reasonable surrogate for atRA binding to BSA, HDL, Apo A-I and RBP4. Considering these proteins’ relative serum and extravascular abundances, and their similar binding affinity to atRA, we predict that serum albumin is an important atRA carrier. Use of floRA in whole tissue tracer studies shows promise but requires further refinement.

## Introduction

The global prevalence of myopia has markedly increased in recent decades [1-3]. Severe myopia substantially enhances the risk of sight-threatening eye conditions such as glaucoma, retinal detachment, and macular degeneration, establishing it as a leading risk factor for blindness worldwide [4]. Myopia usually arises from excessive elongation of the eye along its optical axis and involves remodeling of the scleral extracellular matrix [5]. Clinical and experimental investigations suggest visual input through environmental stimuli is critical for the process that drives the overall axial length of the eye to come into balance with its refractive power, a process known as emmetropization [6]. However, what is not understood is how these changes are processed by the retina and subsequently translated to the changes seen in the sclera, or how disruptions to these processes lead to refractive disorders such as myopia [7].

It is widely recognized that scleral remodeling during myopigenesis involves a retinoscleral signaling cascade, which likely involves multiple signaling molecules, including all-*trans* retinoic acid (atRA) [8-11]. For example, it has been shown that orally delivered exogenous atRA in mice increases the concentration of atRA in several ocular tissue layers (including sclera), induces myopia, and causes scleral remodeling [12], consistent with earlier studies in other species [13-16].

Despite the importance of understanding atRA-linked retinoscleral signaling pathways in myopigenesis, the mechanisms of atRA transport in relevant ocular tissues remain poorly understood. Since atRA is extremely hydrophobic [17], it must bind to one or more carriers for significant transport within aqueous environments [18]. Previous studies identified some of these binding partners in different contexts [19-24], but uncertainty remains about the key binding partner(s) for atRA in the mammalian eye during myopigenesis. Further, quantitative, spatially-resolved measurements of atRA concentrations in tissues are currently infeasible, in part because the weak autofluorescence of atRA limits its direct visualization in tissues using standard fluorescence microscopy. These factors greatly inhibit our understanding of atRA transport.

To better understand the role of atRA in myopigenic retinoscleral signaling, it would be very useful to be able to track the movement of atRA across the sclera and other relevant tissues. There exists a recently developed fluorescent analog of atRA (here denoted as floRA; Figure **1**) [25] that exhibits strong fluorescence when bound by a carrier protein but is non-fluorescent in aqueous environments [25]. This allows one to image the spatial localization of floRA within cells and tissues, positioning it as a useful probe for investigating atRA transport in myopigenesis. To confidently use floRA for this purpose, we must be sure that: (i) atRA and floRA have similar binding affinities to potential carrier proteins, and (ii) it is feasible to quantitatively track the location of a floRA-protein complex in tissue.

**Figure 1.**
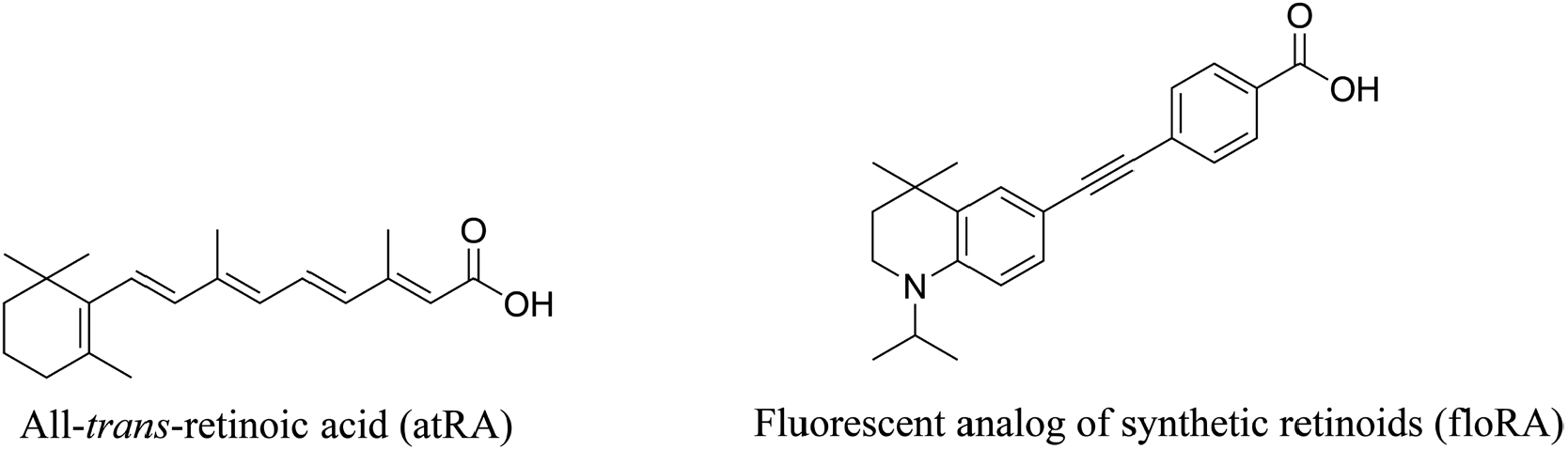
Chemical structures of all-*trans* retinoic acid (atRA) and its fluorescent analog (floRA).

Here we investigated the binding affinity of selected proteins to atRA, focusing on their relevance to retinoscleral transport, and compared atRA and floRA binding to these proteins to evaluate the potential of floRA as a surrogate for atRA in tracer studies. Further, we conducted proof-of-concept experiments to visualize the presence of floRA in relevant ocular tissues from mice.

## Methods

### Reagent specifications and protein production

Commercially available reagents were of reagent grade or better. atRA (≥98% pure, catalog no. R2625, CAS: 302-79-4) was purchased from Sigma-Aldrich (St. Louis, MO, USA) and was used without further purification. floRA (≥98% pure LightOx^TM^ 14, CAS: 198696-03-6) was purchased from LightOx Limited (Newcastle upon Tyne, UK) in lyophilized solid form. Streptavidin Agarose resin (Pierce^TM^ Streptavidin Agarose; catalog no. 20359) was purchased from Thermo Fisher Scientific (Waltham, MA, USA). This resin contains streptavidin recombinant protein crosslinked with 6% beaded agarose with particle size ranging from 45-165 μm. Sources of purchased proteins, and their concentrations as received, are given in Table **S1**.

#### Cellular retinoic acid-binding protein 2 (CRABP2)

Mouse CRABP2 protein was expressed in One Shot^®^ BL21(DE3) *E. coli* (Thermo Fisher Scientific, Waltham, MA, USA) according to the manufacturer’s instructions, using a plasmid custom synthesized by GenScript (NJ, USA). Protein purification was performed using Cytiva Glutathione Sepharose^TM^ 4B (MilliporeSigma, MA, USA), and the GST tag was removed by PreScission Protease (GenScript, NJ, USA). Buffer exchange of purified mouse CRABP2 used Amicon® Ultra-15 Centrifugal Filter 3K Devices (MilliporeSigma, MA, USA). Mouse CRABP2 protein concentration was determined from absorbance measurements at 280 nm using a calculated ε value of 19,480 M^-1^cm^-1^ [26]. Purity was assessed by subjecting 1-3 μg of purified protein to SDS-PAGE followed by determination of band intensity after Coomassie blue staining. The purified protein was then stored at -80°C in 1x PBS buffer solution.

#### Cellular retinol-binding protein 1 (RBP1)

Mouse RBP1 was prepared according to the protocol reported in [27, 28]. In brief, Mouse RBP1 was expressed in BL 21 *E. coli* purchased from Sigma-Aldrich, utilizing plasmids sourced from Genecopia (Rockville, MD, USA) as per the manufacturer’s instructions. Purification procedures involved the use of a GE Healthcare GST bulk kit (GE Healthcare, Pittsburgh, PA, USA). Following expression, the GST tag was cleaved using Promega ProTEV protease (Promega, Madison, WI, USA). Subsequently, the removal of the protease was facilitated by GE Healthcare Ni resin. To ensure purity, the protein solution underwent a second pass through the GST column to separate the GST tag from the purified protein. The purified protein was then dialyzed and stored at -80°C in 20 mM KH_2_PO_4_ and 100 mM KCl buffer.

### Fluorescence and UV-Vis spectroscopic measurements

All spectra were acquired at room temperature. Fluorescence spectroscopic measurements were acquired with the sample solutions of volume ∼700 μL in a 1.0 cm pathlength fluorescence quartz cuvette (FireflySci, Inc., NY, USA). Fluorescence spectra were recorded using a Horiba Jobin Yvon FL3-21 Fluorescence Spectrophotometer (Horiba Ltd., Kyoto, Japan) equipped with a 450-W Xe lamp, at a scan rate of 120 nm/min. The spectra were processed with FluorEssence^TM^ (Horiba Ltd., Kyoto, Japan) software and plotted and analyzed using MATLAB 2023a (The MathWorks, Natick, MA). In the fluorescence quenching experiments, the excitation wavelength was 280 nm, and the emission spectra were obtained in a wavelength range of 300-450 nm (protein-atRA binding) and 300-550 nm (protein-floRA binding). In fluorescence potentiation experiments with floRA, the excitation wavelength was 340 nm, and the emission spectra were obtained in a wavelength range of 400-600 nm.

UV-Vis absorbance spectra were obtained using an Evolution^TM^ 220 UV-Vis spectrophotometer (Thermo Fisher Scientific, Waltham, MA, USA) with the sample solutions of volume ∼700 μL in a 1.0 cm pathlength quartz cuvette (FireflySci, Inc., NY, USA). Prior to recording the spectra of the sample solutions, a baseline measurement was conducted using the same PBS buffer solution employed throughout the experiment. This baseline correction was then automatically applied to each spectrum by using default software settings. The spectra were processed with INSIGHT^TM^ software (Thermo Fisher Scientific, Waltham, MA, USA), plotted, and further analyzed with MATLAB. The measurements were conducted across the UV-Vis region. However, we here focus on presenting spectra over the range 250–550 nm, since no significant spectral features were observed outside this range.

For comparative binding experiments with atRA, Apo A-I and biotinylated BSA, fluorescent spectra were recorded utilizing a FlexStation3 (Molecular Devices, LLC., CA, USA) microplate reader, processed with SoftMax® Pro 5.4.6 (Molecular Devices, LLC., CA, USA) software, plotted, and analyzed using MATLAB. Each experiment utilized 40 μL of sample solution.

### Determination of protein-atRA and protein-floRA association constants and stoichiometry from fluorescence quenching data

To determine the association constants of protein-atRA/floRA binding, we utilized the quenching of the proteins’ intrinsic tryptophan fluorescence through complexation between the carrier protein and atRA/floRA (Figure **S1**). The protein concentrations ranged from 1-6 μM and the amounts of atRA/floRA titrated ranged from 0-25 μM. Full details are provided in Supplemental Table **S2**. Upon addition of atRA/floRA to the protein solution, the sample was mixed well and incubated on ice for 15-20 minutes before measurement of fluorescence spectra. The results from fluorescence measurements were fitted to standard models (Stern-Volmer, modified Stern-Volmer [21-23, 29, 30], quadratic [24, 27, 28]) to estimate the association constants of retinoid–protein complexation, as described in detail in the Supplemental material. In the main text we report values of the association constant based on quadratic fitting, since the quadratic model can deal with nonlinear trends in fluorescence quenching data at high concentrations. In Table **1**, we show association constant values based on all fitting approaches to allow comparison with previous literature reports. Detailed results from other fitting approaches are shown in the Supplemental Material.

**Table 1.** Association constant values (K_a_, mean ± SD) estimated from fluorescence quenching experiments with atRA and floRA, as obtained by different fitting approaches (see Methods). Abbreviations: SV = Stern-Volmer; mSV = modified Stern-Volmer. Dashes indicate conditions where measurements/K_a_ estimations were not made.

| Protein | Ligand | Association constant |  |  |  |
| --- | --- | --- | --- | --- | --- |
|  |  | SV model | mSV model |  | Quadratic model |
| | | $K_a (\times 10^5 \text{ M}^{-1})$ | $K_a (\times 10^5 \text{ M}^{-1})$ | n | $K_a (\times 10^5 \text{ M}^{-1})$ |
| BSA | atRA | $1.6 \pm 0.2$ | $1.4 \pm 0.4$ | $1.1 \pm 0.3$ | $2.4 \pm 0.3$ |
| | floRA | $1.3 \pm 0.1$ | $1.0 \pm 0.4$ | $1.3 \pm 0.4$ | $2.1 \pm 0.4$ |
| HDL | atRA | $6.4 \pm 0.3$ | $6.9 \pm 0.9$ | $1.0 \pm 0.1$ | $10.1 \pm 5.4$ |
| | floRA | $5.9 \pm 0.2$ | $3.4 \pm 0.0$ | $1.4 \pm 0.0$ | $13.2 \pm 1.1$ |
| Apo A-I | atRA | $1.2 \pm 0.2$ | $1.3 \pm 0.4$ | $1.0 \pm 0.3$ | $6.3 \pm 0.3$ |
| | floRA | $1.7 \pm 0.5$ | $1.5 \pm 0.5$ | $1.1 \pm 0.2$ | $7.9 \pm 0.2$ |
| RBP4 | atRA | $1.5 \pm 0.2$ | $1.7 \pm 0.8$ | $1.0 \pm 0.3$ | $6.6 \pm 0.2$ |
| | floRA | $0.3 \pm 0.0$ | $0.6 \pm 0.1$ | $0.7 \pm 0.1$ | $4.3 \pm 0.2$ |
| CRABP2 | atRA | - | - | - | $784 \pm 634$ |
| | floRA | - | - | - | $669 \pm 92$ |
| RBP1 | atRA | - | - | - | - |
| | floRA | $1.5 \pm 0.3$ | $1.4 \pm 1.3$ | $1.2 \pm 0.4$ | $7.9 \pm 0.4$ |

### Determination of binding constants from fluorescence potentiation experiments with floRA

Fluorescence emission of floRA in an aqueous solution is severely quenched, but this fluorescence is increased in a concentration-dependent manner when floRA is bound to a hydrophobic binding pocket of a protein in an aqueous solution [25]. We exploited this behavior and implemented an experimental approach, denoted as “fluorescence potentiation experiments” to estimate the floRA-protein association constant. In these experiments, the floRA concentration was fixed, while the protein concentration was gradually increased causing the fluorescence emission intensity to reach a plateau at a certain protein concentration threshold [25]. The floRA concentration ranged from 0.025-0.05 μM, and the protein concentrations titrated into the floRA solution ranged from 0-35 μM. Full details are provided in Supplemental Table **S3**. Upon the addition of floRA to the protein solution, the sample was mixed well followed by incubation on ice for 15-20 minutes before fluorescence measurement. We fitted the Hill Equation (taking the Hill coefficient as unity, i.e. ignoring the potential effect of cooperativity in protein-ligand binding) to the fluorescence data to evaluate the binding constant [31] using the ‘fit’ function in MATLAB. The Hill equation is expressed in relation to the fluorescence data as:

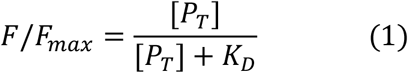

where [P_T_] is the total protein concentration (μM), *F*_*max*_ represents the fluorescence signal obtained at the spectral maxima once the saturation plateau is attained and *F* is the fluorescence intensity at protein concentration [*P*_*T*_]. *K*_*D*_ represents the dissociation constant (estimated from the fitting), the inverse of which is taken as the association constant *K*_*a*_ (M^-1^).

### Protein-ligand complex isolation protocol

A mixture of 1 mM floRA in ethanol (0.08 mL) and 0.1 mM BSA in PBS (1.4 mL) was passed through a PD-10 desalting column (freshly equilibrated with PBS buffer), with an additional 1.0 mL of PBS added concurrently. The initial eluate (∼2.5 mL) was discarded, and the BSA-floRA complex was isolated by adding a total of 4.0 mL of PBS in 0.5 mL increments, with eluate collected into 8 separate Eppendorf tubes. UV-Vis absorbance revealed the presence of BSA-floRA in the first three eluate fractions, which were combined and used directly for incubation with mouse scleral tissue, as described in the section titled ‘Visualization of floRA in ocular tissues’ below.

### Competitive binding experiments with atRA, Apo A-I and Biotinylated BSA

Polypropylene chromatography columns were packed with 2 mL of streptavidin-agarose resin, followed by equilibration with PBS (2 × 10 mL). Biotinylated BSA (50 μM, 1 mL) in PBS was passed through the column. The eluate was collected and passed through the column twice more, after which no UV-visible absorbance was detected in the eluate indicating complete adsorption of the biotinylated BSA by the streptavidin resin. A solution of atRA (50 μM, 1 mL) in PBS and ≤ 5% ethanol (EtOH) was then passed through the column. The eluate was then collected and analyzed, revealing no detectable absorbance for either biotinylated BSA or atRA, indicating successful BSA-atRA binding.

To investigate whether atRA selectively favors one potential carrier over another, a solution of 50 μM Apo A-I and 50 μM atRA was passed through a streptavidin-biotinylated BSA column (prepared as above). The eluate was then collected and circulated through the column four times, and the UV-Vis absorbance of each eluate was recorded. Subsequently, ∼10-15% ethanol was added to the fourth eluate. This combined solution was passed through the column and collected for spectroscopic measurements.

### Visualization of floRA in ocular tissues

All animal procedures adhered to the ARVO Statement for the Use of Animals in Ophthalmic and Vision Research, and to a protocol approved by the Institutional Animal Care and Use Committee at Emory University. Male and female C57BL/6J mice were procured from Jackson Laboratory (Bar Harbor, Maine, USA) and housed in the animal facility at Emory University on a 12-hour light-dark cycle. Mice approximately 4-5 weeks of age were euthanized by cervical dislocation. Their eyes were then enucleated, placed in 0.1 M (1x) phosphate-buffered saline (PBS) (Thermo Fisher Scientific, Waltham, MA), dissected to carefully remove extraocular muscle, fat, and conjunctiva, and hemisected at the corneoscleral junction. The anterior segment and intraocular tissues were removed to leave scleral shells with adherent choroid. Each posterior shell was cut into quadrants, which were further cut into pie-shaped wedges. These were incubated for various durations (5 min, 10 min, 30 min, 60 min, 90 min and 180 min) at 4°C in a BSA-floRA solution (approximately 50 μM floRA-BSA complex in PBS and ≤ 5% EtOH). Prior to incubation, this solution underwent a protein-ligand complex isolation protocol as described above.

Tissue wedges were embedded in OCT compound (Fisher HealthCare, Houston, TX), snap-frozen and stored at -80°C. Sections, 10 μm thick, were cut on a cryostat and collected on glass slides. To reduce diffusion of floRA out of the tissue sections, solution changes and washes were eliminated. Specifically, to stain nuclei, we prepared a mixture of ProLong^TM^ Gold media and Sytox^TM^ Orange (both from Thermo Fisher Scientific, Waltham, MA, diluted 100:1 respectively) that was vortexed briefly, then centrifuged. After collected sections were allowed to dry for 10 min, slides were mounted using the above mixture and cover-slipped, omitting the permeabilization step and washes. The slides were allowed to cure overnight at 4°C, then imaged with a Leica DM6 fluorescent microscope (Leica Microsystems, Wetzlar, Germany) equipped with the appropriate filter cubes. Since floRA was found to be highly sensitive to bleaching and fading, illumination was strictly minimized in terms of exposure time and intensity, and imaging for each experiment was accomplished within 2h. Brightfield and fluorescent (floRA and Sytox^TM^ Orange channels) images were acquired (16 bit, 2048 × 2048 pixels, 6.2 pixels/micron) and stored for later analysis.

Fluorescent images showed significant local spatial heterogeneity in floRA intensity in both the sclera and choroid, making extraction of reliable spatially-resolved fluorescence intensity profiles infeasible. Instead, we quantified average floRA fluorescence intensity within a large region of the sclera at different time points, thereby seeking to reduce the effects of spatial heterogeneity by spatial averaging. Specifically, by simultaneous inspection of paired brightfield and floRA fluorescence images in Fiji v. 1.54k [32], we manually drew curves along the outer scleral margin and the scleral-choroidal interface. We then connected these curves by straight lines across the sclera (Figure **S2**), to create a quadrilaterally-shaped scleral region. The mean fluorescent intensity (after correction for background fluorescence) within this region and the area of this region were obtained in Fiji. This was repeated on 4 sections per timepoint (technical replicates) for two eyes (biological replicates). The area of the analyzed scleral region varied somewhat from one section to another, but was typically 8700 square microns, and scleral thickness was typically ∼40 microns.

## Results

### Overview of atRA/floRA protein binding experiments

The suitability of floRA as a fluorescent atRA analog in tracer studies was studied by characterizing the binding of both atRA and floRA to the following carrier proteins using protein-dependent changes in fluorescence spectra: bovine serum albumin (BSA), high-density lipoprotein (HDL), apolipoprotein A-I (Apo A-I, a part of the high-density lipoprotein (HDL) complex [33]), and retinol binding protein 4 (RBP4). We also used cellular retinoic acid-binding protein 2 (CRABP2) and cellular retinol-binding protein 1 (RBP1) as positive and negative controls, respectively, due to their known binding affinities with atRA [24, 27, 34, 35]. We chose BSA and HDL because they are potential carriers of atRA through the extracellular space, including within the bloodstream. Apo A-I has been implicated as an atRA binding protein in the eye in myopia [19], while RBP4, a known chaperone for all-*trans*-retinol [36-38], which is known to bind retinol and atRA with similar binding affinity [38, 39] may facilitate atRA transport before cellular uptake. Uptake of retinoic acid has been observed in pigment epithelial cells *in vitro* through dissociation of retinoic acid-RBP complex [40].

All the above proteins have intrinsic tryptophan fluorescence that can be quenched upon atRA binding to the protein [19, 23, 24]. We made use of this phenomenon to evaluate the binding affinities of atRA to these potential carrier proteins. Further, because atRA can bind to multiple carrier proteins [19-24], we also investigated atRA’s propensity to switch from one protein carrier to another.

### Control experiments with CRABP2 and RBP1

Previous research has quantified the strong affinity of both atRA and floRA for CRABP2 [24, 25, 35]. Thus, we chose CRABP2 as a positive control for our fluorometric titration experiments. The expected quenching of CRABP2 tryptophan fluorescence was observed upon the addition of atRA or floRA (Figure **S3**), with no noticeable alterations detected in other spectroscopic parameters. We also observed an additional emission peak around 463 nm (upon excitation at 280 nm, Figure **S4**) due to the intrinsic fluorescence of the floRA in the presence of CRABP2 (Figure **S3C**). Association constants of atRA and floRA with CRABP2 determined using these fluorescence signals were closely comparable (Tables **1****, S4**) and aligned well with the lower range of values documented in the literature (for atRA-CRABP2, 700-5000 × 10^5^ M^-1^ [24, 34, 35]; for floRA-CRABP2, 300 × 10^5^ M^-1^ [25]).

Conversely, previous studies have shown that atRA does not bind to RBP1 [27, 28], suggesting its utility as a negative control. In fluorescence quenching experiments, we found that RBP1 tryptophan emission intensity did not change as a function of atRA concentration, confirming its expected lack of binding (Figure **S5A-S5B**). However, similar experiments using floRA instead of atRA showed quenching of RBP1 fluorescence by floRA in a concentration-dependent manner (Figure **S5C-S5F**), indicating dissimilarities between atRA and floRA when interacting with RBP1. Quenching of RBP1 fluorescence by floRA allowed us to estimate an apparent association constant of 7.9 ± 0.4 × 10^5^ M^-1^ (Table **1**).

### atRA and floRA show comparable binding affinities to BSA, HDL and Apo A-I

The binding of atRA to BSA was similarly quantified using BSA’s intrinsic tryptophan fluorescence quenching (λ_em_ = 336 nm, λ_ex_ = 280 nm) upon ligand exposure [21-23, 41] (Figures **2A****, S1, S6A-S6B**). atRA alone is non-fluorescent in water and the addition of atRA to BSA induces no other discernible spectroscopic alterations (Figure **2A-2B**, Figure **S7A-S7B**). The addition of floRA to BSA caused a similar loss of BSA tryptophan fluorescence emission in a concentration-dependent manner (Figure **2C-2D**, Figure **S7C-S7D**), along with the appearance of an additional emission peak at 461 nm (λ_ex_ = 280 nm) due to the intrinsic fluorescence of the floRA in the presence of BSA (Figure **2C**). floRA alone, in the absence of BSA, remained non-fluorescent in an aqueous environment (Figure **S8**).

**Figure 2.**
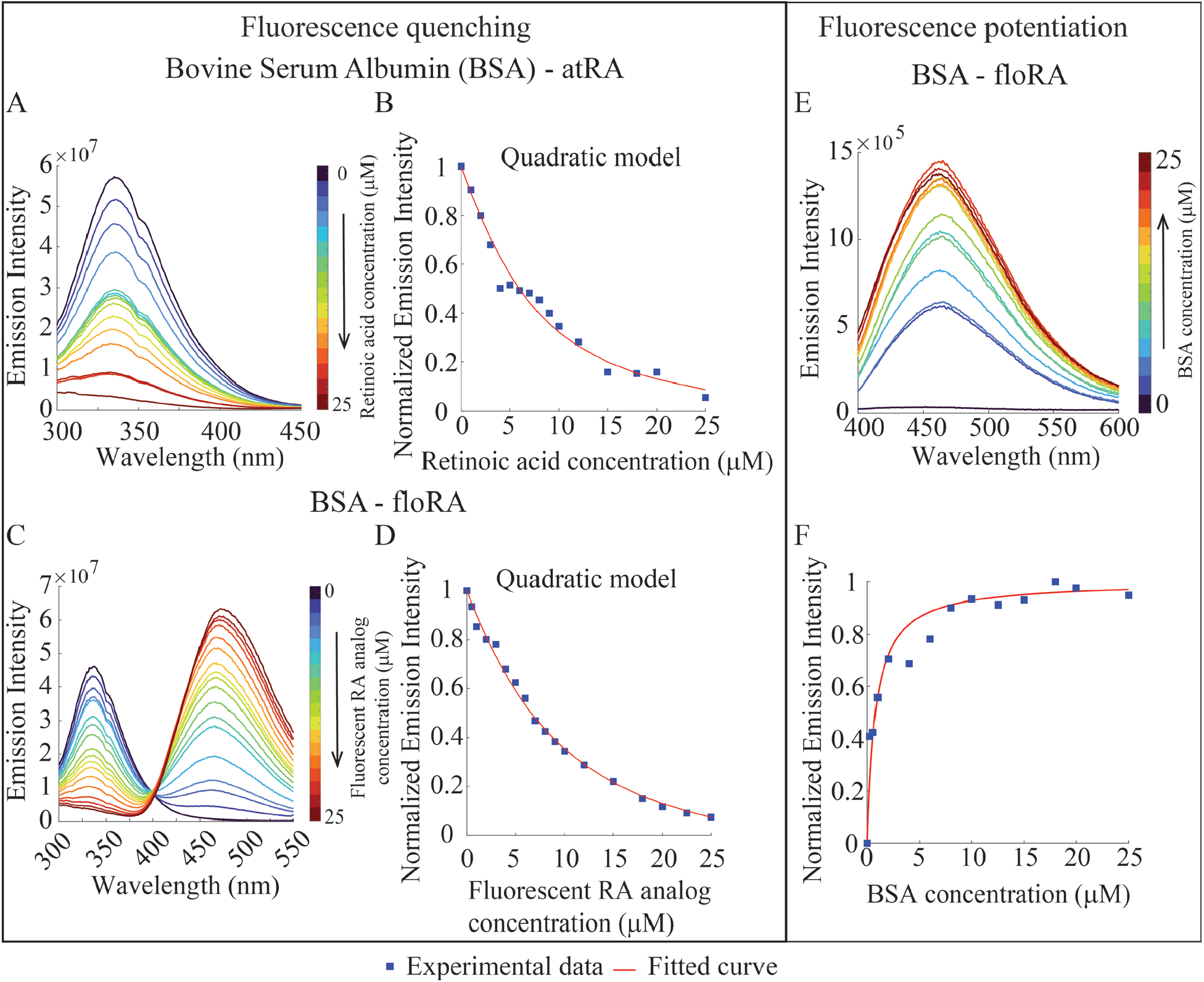
Fluorometric investigations of BSA-retinoic acid binding. **(A, C)** Fluorescence emission spectra of 6 μM BSA in the presence of atRA or floRA, respectively (0-25 μM). The quenching of BSA fluorescence (336 nm) with increasing concentrations of atRA or floRA is evident. **(B, D)** Quadratic model fits to the peak emission intensities at 336 nm from the spectra in panels **A** and **C. (E)** Fluorescent emission spectra of floRA (0.05 μM) in the presence of BSA (0-25 μM); increase in peak intensity indicates increasing amounts of floRA bound by the protein. **(F)** Hill curve fit of fluorescence intensity (461 nm) in panel **E**. Panels **A-F** show typical results from 3 technical replicates.

Using tryptophan fluorescence data, the association constants for BSA-atRA and BSA-floRA binding were found to be 2.4 ± 0.3 × 10^5^ M^-1^ and 2.1 ± 0.4 × 10^5^ M^-1^, respectively (Table **1**), consistent with previous reports for albumin-atRA binding (3-23 × 10^5^ M^-1^ [21-23, 42]). The concentration-dependent appearance of floRA fluorescence in complexation with BSA (Figure **2E-2F**) provided an estimated association constant of 11.8 ± 1.2 × 10^5^ M^-1^ (Table **S4**). While somewhat different than the tryptophan-derived value, the binding affinities of atRA and floRA to BSA are of the same order of magnitude, enhancing confidence in floRA’s potential role as a probe for investigating BSA-mediated atRA transport.

Previous work has indicated that Apo A-I, a major constituent of HDL, may be an important carrier of atRA in myopigenesis [19]. Both tryptophan fluorescence quenching and floRA fluorescence potentiation were observed upon the addition of atRA and floRA to HDL and Apo A-I in a similar manner to BSA (Figures **3** and **4**; see also supporting data in Figures **S9, S10**, and Tables **S2**-**S4**). As was the case with BSA, fitting of the fluorescence quenching data showed that HDL-atRA (10.1 ± 5.4 × 10^5^ M^-1^) and HDL-floRA (13.2 ± 1.1 × 10^5^ M^-1^) association constants were comparable (Table **1**), as was the value determined using HDL-floRA fluorescence potentiation (10.3 ± 3.5 × 10^5^ M^-1^, Table **S4**). Of note, since the HDL solution we used, contains multiple proteins including Apo A-I and Apo A-II and their relative fraction was unknown, we determined the effective protein concentration within HDL by treating this concentration as a free parameter in the quadratic model fitting, allowing us to convert the known HDL concentration (in terms of mg/ml) into μM. The details of this analysis are provided in the Supplemental material.

**Figure 3.**
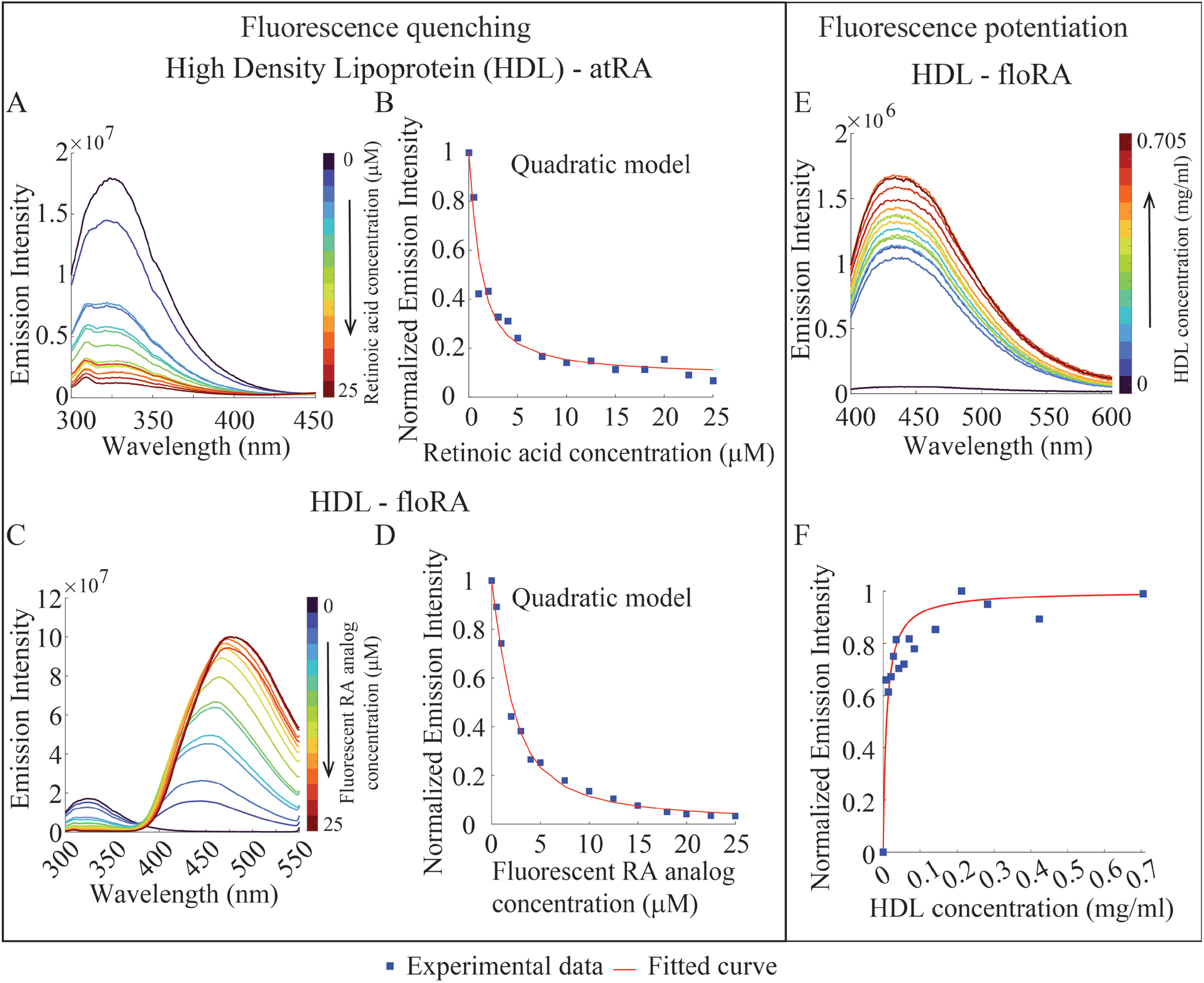
Fluorometric investigations of HDL-retinoic acid binding. **(A, C)** Fluorescence emission spectra of HDL (0.028 mg/mL) in the presence of atRA or floRA, respectively (0-25 μM). The quenching of HDL fluorescence (321 nm) with increasing concentrations of atRA or floRA is evident. **(B, D)** Quadratic model fits to the peak emission intensities at 321 nm from the spectra in panels **A** and **C. (E)** Fluorescent emission spectra of floRA (0.05 μM) in the presence of HDL (0-0.7 mg/mL); the increase in peak intensity indicates increasing amounts of floRA bound to HDL. **(F)** Hill curve fit of fluorescence intensity (437 nm) in panel **E**. Panels **A**-**F** show typical results from 3 technical replicates.

**Figure 4.**
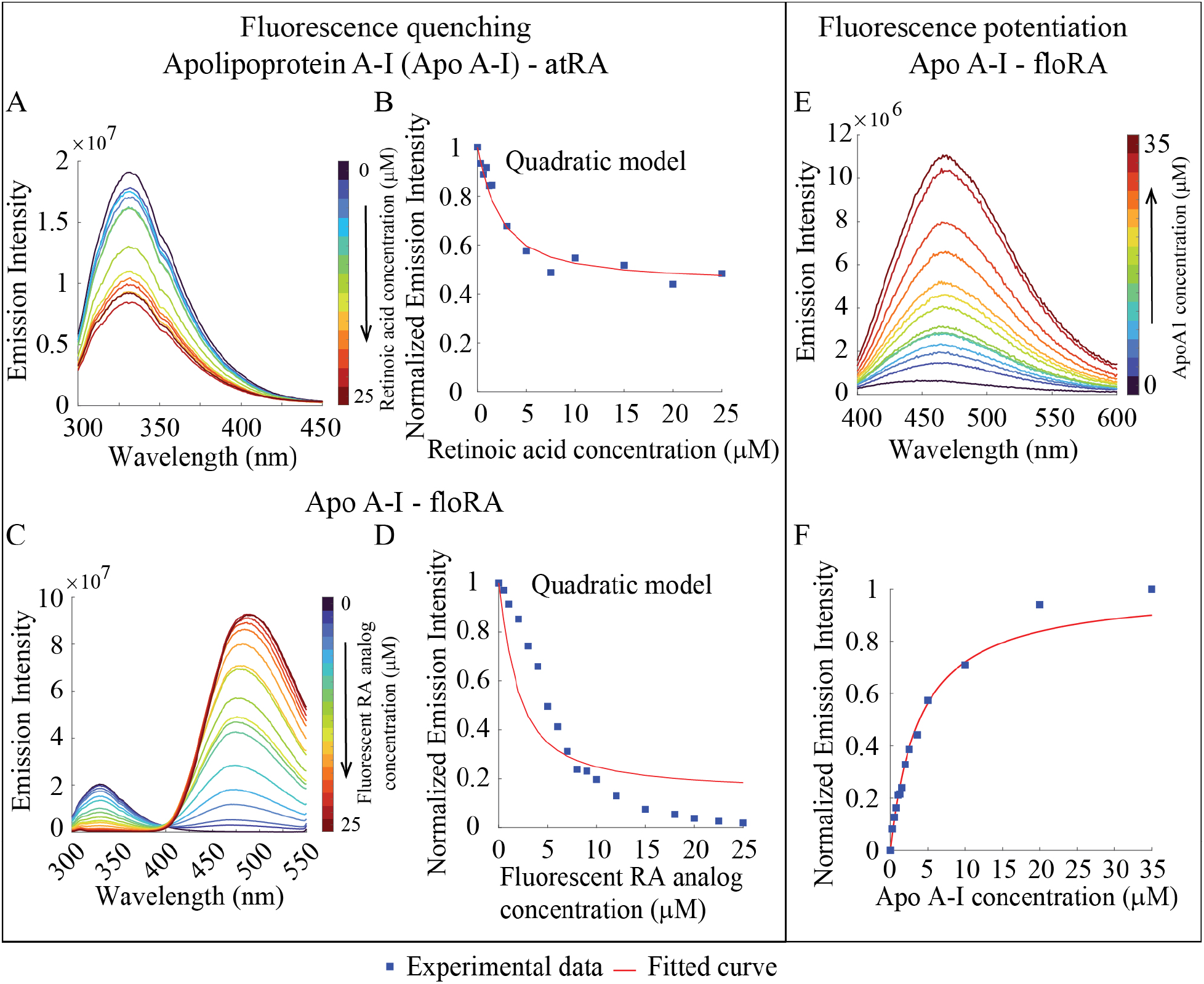
Fluorometric investigations of Apo A-I-retinoic acid binding. **(A, C)** Fluorescence emission spectra of Apo A-I (1.2 μM) in the presence of atRA or floRA, respectively (0-25 μM). The quenching of Apo A-I fluorescence (332 nm) with increasing concentrations of atRA or floRA is evident. **(B, D)** Quadratic model fits to the peak emission intensities at 332 nm from the spectra in panels **A** and **C. (E)** Fluorescent emission spectra of floRA (0.05 μM) in the presence of Apo A-I (0-35 μM); the increase in peak intensity indicates increasing amounts of floRA bound to the protein. **(F)** Hill curve fit of the fluorescence intensity (467 nm) in panel **E**. Panels **A**-**F** show typical results from 3 technical replicates.

Apo A-I binding led to generally similar fluorescence behavior as HDL [19], but we observed disagreement between data and quadratic model fitting in Figure **4D**. The discrepancy may stem from the molecular interactions between floRA and Apo A-I affecting the energy transfer during the tryptophan fluorescence quenching. While all data fitting models assume static quenching, in which the quencher forms a non-fluorescent ground state complex, the existence of dynamic quenching cannot be ruled out. Further, it is documented that Apo A-I undergoes self-assembly and form Apo A-I globular condensates [43]. Previous studies have shown that protein aggregation can induce self-quenching of fluorescence suggesting that the aggregation induced self-quenching could also contribute to the observed discrepancy [44, 45]. Therefore, although the fitted association constants for Apo A-I-atRA binding (6.3 ± 0.3 × 10^5^ M^-1^) and Apo A-I-floRA binding (7.9 ± 0.2 × 10^5^ M^-1^) were similar (Table **1**), there may be some differences in floRA-Apo A-I binding vs. atRA-Apo A-I binding. The binding constant obtained from fluorescence potentiation of floRA upon Apo A-I binding (2.5 ± 0.1 × 10^5^ M^-1^, Figure **4E-4F** and Table **S4**) was again similar to that obtained by tryptophan fluorescence quenching.

### atRA and floRA binding to RBP4

It is known that retinol (ROL) circulating in the bloodstream is transported by retinol-binding protein 4 (RBP4) [46-49]. We characterized RBP4 binding to atRA [38, 39] and floRA by examining the quenching of RBP4 tryptophan emission upon binding with atRA and floRA (Figure **5A-5D**, Figure **S11**, Table **S2**). The derived association constants for RBP4-atRA and RBP4-floRA were again similar, 6.6 ± 0.2 × 10^5^ M^-1^ and 4.3 ± 0.2 × 10^5^ M^-1^, respectively (Table **1**). Interestingly, even with higher concentrations of the quencher floRA, the peak tryptophan fluorescent emission intensity was reduced by only ∼20% (Figure **5D**), suggesting some differences in the binding geometries of atRA vs. floRA to RBP4. This was supported by the lack of saturated fluorescence turn-on of floRA by interaction with RBP4 (Figure **5E-5F**).

**Figure 5.**
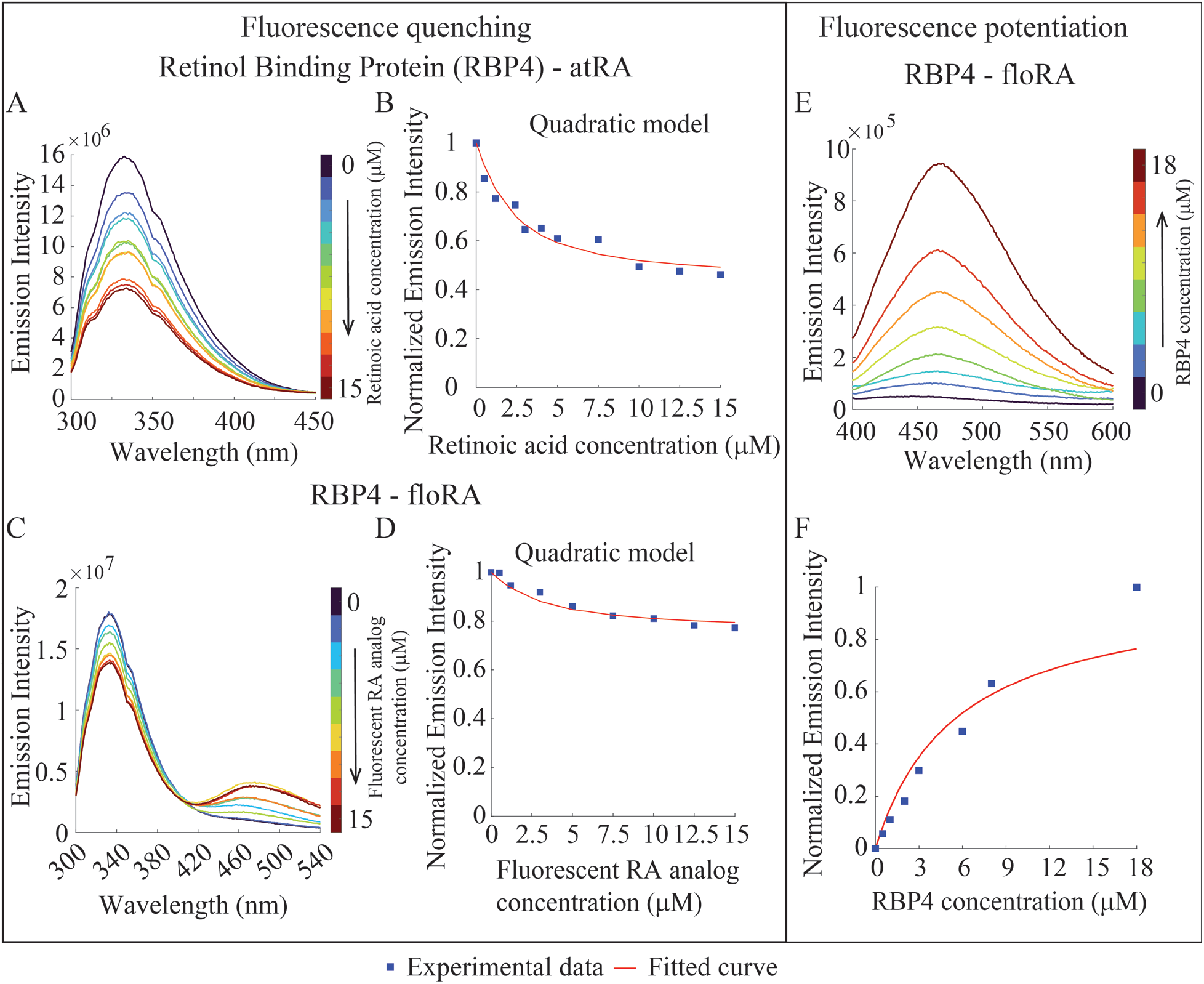
Fluorometric investigations of RBP4-retinoic acid binding. **(A, C)** Fluorescence emission spectra of RBP4 (1.2 μM) in the presence of atRA or floRA, respectively (0-15 μM). The quenching of RBP4 fluorescence (334 nm) with increasing concentrations of atRA or floRA is evident. **(B, D)** Quadratic model fits to the peak emission intensities at 334 nm from the spectra in panels **A** and **C. (E)** Fluorescent emission spectra of floRA (0.025 μM) RBP4 (0-18 μM); the increase in peak intensity indicates increasing amounts of RBP4 bound to the protein. **(F)** Hill curve fit of the fluorescence intensity (466 nm) in panel **E**. Panels **A**-**F** show typical results from 3 technical replicates.

### Competition between immobilized and solution-phase binding proteins for atRA

We explored whether atRA would selectively bind to one putative carrier over another in a competition between a surface-immobilized protein and one in solution, behavior that is relevant to the physiological environment where multiple potential carriers are present, some of which may be adherent to extracellular matrix or cells. Biotinylated BSA, immobilized on a column of streptavidin beads, was shown to be able to efficiently capture atRA from solution (Figure **6A**) and to compete with soluble Apo A-I for that same analyte (Figure **6B, C**). After several passages of the Apo A-I-atRA mixture through the immobilized BSA, atRA was completely retained on the column and was not dislodged by an additional wash with 10-15% ethanol (Figure **S12**). Control experiments showed no binding of atRA to the bead material (Figure **S12D-S12F**). Since the solution-phase association constants of BSA and Apo A-I with atRA are very similar (Table **1**), this experiment may highlight the difference between binding to an immobilized protein binding partner (defined by an adsorption coefficient, *K*_ADS_) and a solution-phase binding partner (defined by association constant, *K*_a_). For example, in a well-characterized example of carbohydrate-lectin binding, *K*_ADS_ was found to exceed *K*_a_ by a factor of approximately 350 [50].

**Figure 6.**
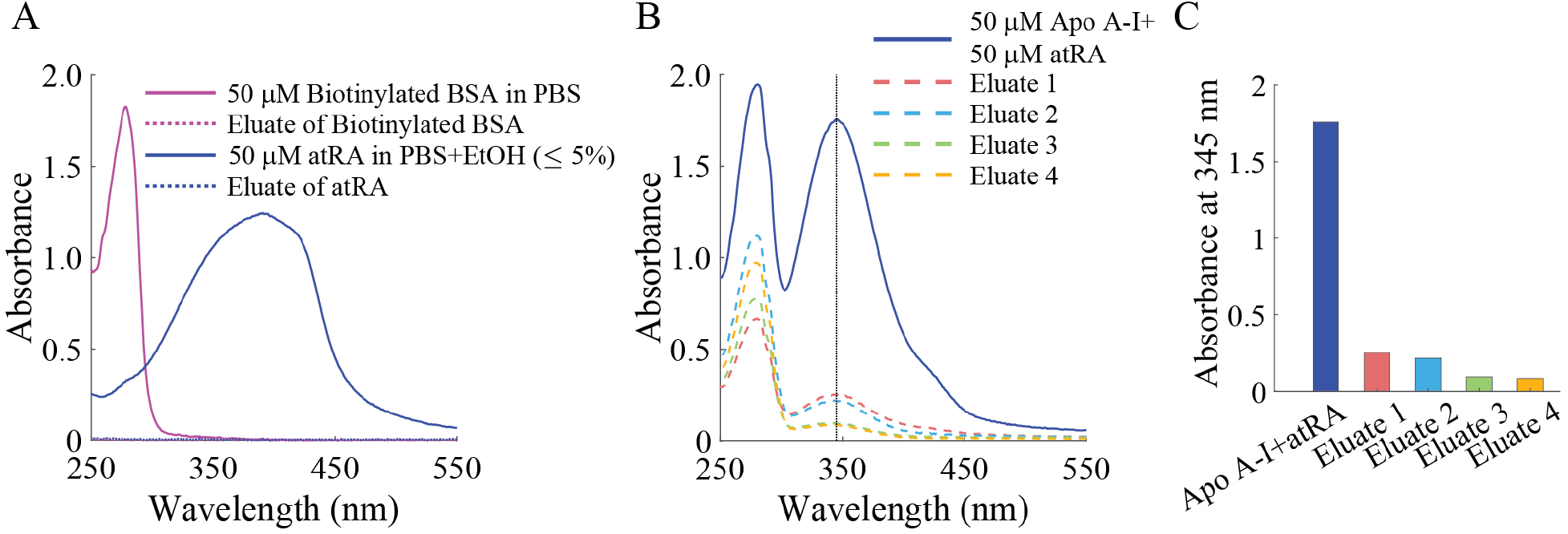
atRA shows preferential binding to immobilized BSA vs. soluble Apo A-I. **(A)** UV-Vis absorbance spectra of separate solutions of biotinylated BSA (50 μM) and atRA (50 μM), and eluates after sequential passage of these solutions through streptavidin resin, showing efficient capture of biotinylated BSA by the resin and atRA by the immobilized BSA. (Dotted curves are difficult to visualize as they partially overlap the horizontal axis.) **(B)** UV-Vis absorbance spectra of Apo A-I-atRA mixture (solid curve) and eluates after passing through a column bearing immobilized BSA. (**C**) Absorbance intensities of Apo A-I-atRA mixture and subsequent eluates at 345 nm from panel **B**. Panels **A**-**C** show typical results from 3 technical replicates.

### floRA can be visualized within *post mortem* ocular tissues, with some limitations

To further investigate floRA’s potential utility as a fluorescent tracer within tissue, we exposed mouse choroid/sclera samples to a 50 μM floRA-BSA mixture, allowing the mixture to passively diffuse into the tissue for various lengths of time. Since neither unbound floRA nor BSA fluoresce at 450-490 nm in aqueous solution when excited at 340-380 nm, any fluorescence we observe in the tissue (above background levels) indicates the presence of protein-floRA complexes.

Generally speaking, the amount of fluorescent labeling in both choroid and sclera increased with increasing incubation duration (Figure **7A**). However, this general trend was non-monotonic, e.g. there was markedly less scleral fluorescence at 30 minutes than at 10 minutes in the example shown (Figure **7A**). In a homogeneous material being passively labeled by a diffusive process, one would expect maximum fluorescence at the free surfaces (external margin of the sclera and internal margin of the choroid). However, this was not the case in these experiments: we frequently observed elevated fluorescence in the vicinity of the choroidal-scleral interface, suggesting that the floRA-atRA complex may have rapidly penetrated into the suprachoroidal space in this *post mortem* preparation. We also observed significant spatial heterogeneity, with alternating bright and dark regions within the sclera. Although some of this pattern could be due to tissue deformation (bending) during histological processing, an alternative explanation is that floRA was sequestering within scleral cells. In fact, co-staining for nuclei showed nuclei residing within bright regions of floRA labeling, consistent with this hypothesis (Figure **7B**). It is of interest that a previous study also observed punctate fluorescent structures in HaCaT cells treated with floRA, suggesting the localization of floRA in nuclei and cytoplasmic lipid vesicles [25].

**Figure 7.**
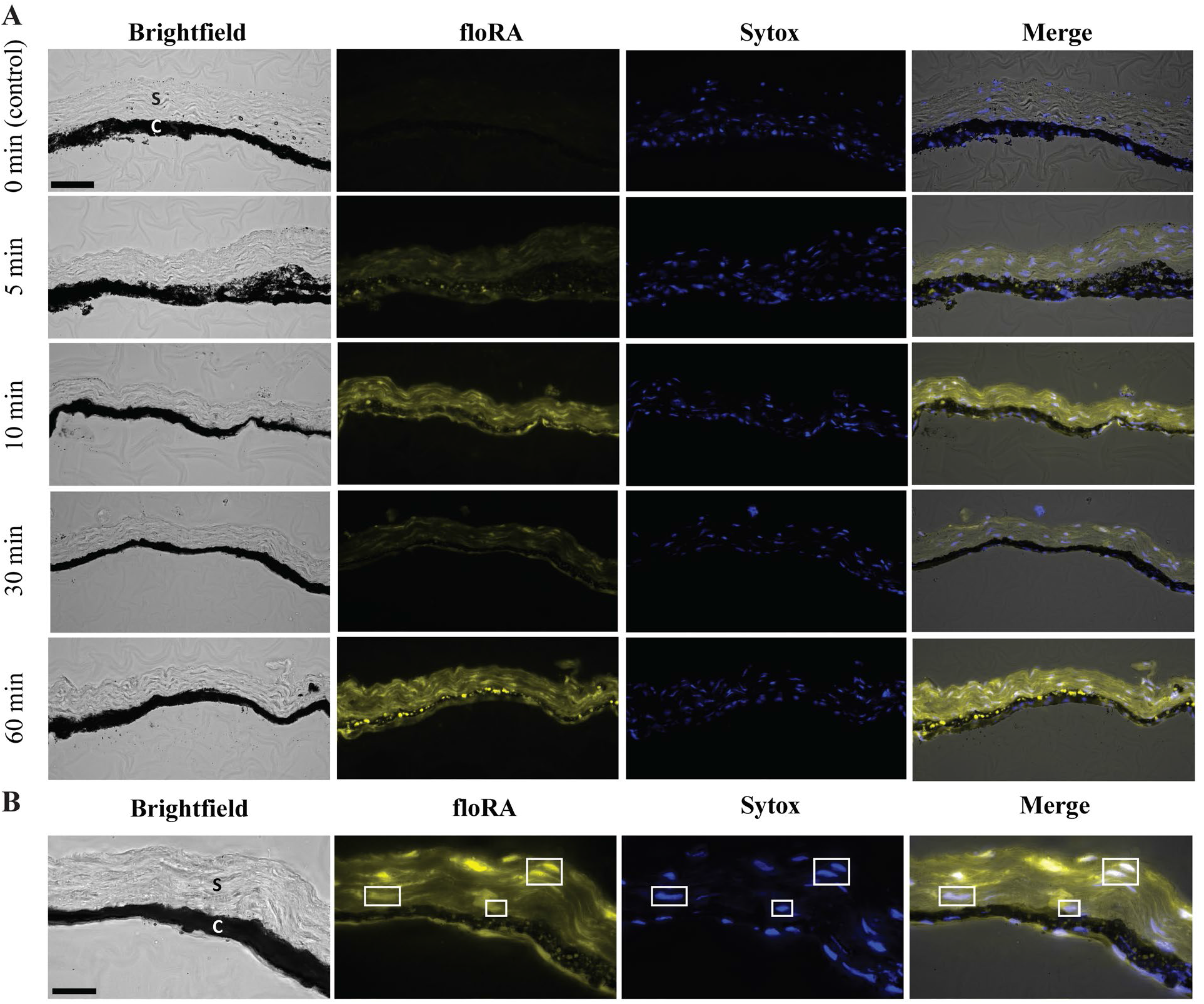
Incubation of mouse choroid/sclera wedges in a solution of floRA and BSA shows floRA penetration into the tissue. **(A)** Tissue wedges incubated in floRA-BSA for various durations, including a 0 minute control “incubation”. We show only micrographs up to 60 minutes, since images at later times showed no additional significant features beyond those in this igure. C = choroid, S = sclera, Sytox = nuclear stain. Scale bar 50 μm. **(B)** Visualization of floRA and nuclei showing co-localization, with selected regions highlighted in white boxes. Abbreviations as in panel **A**. Scale bar 20 μm.

We attempted to fit the quantitative fluorescence data to a model of unsteady diffusion in a homogenous material [51], accounting for the fact that the different scleral samples had different scleral thicknesses and taking the fitted quantity as the spatially averaged concentration of the fluorophore over the entire thickness of the sclera. The fitted quantities were the diffusivity of the floRA-BSA complex within the sclera and the limiting concentration of floRA-BSA in the sclera, i.e. the concentration as incubation time tended to infinity. We assumed that the measured fluorescence was linearly proportional to floRA-BSA concentration. Generally speaking, the fit was poor, e.g. taking different starting guesses for the values of the fitted parameters in the nonlinear fitting process produced significantly different outcomes. We conclude that the distribution of fluorescence in the tissue was not well-described by a passive diffusion process in a homogeneous medium.

## Discussion

We surveyed the binding of several putative carrier proteins with atRA or its synthetic fluorescent analog floRA [25] to evaluate the potential of using floRA as a surrogate tracer for atRA in studies of myopigenesis, as well as to identify possible atRA carrier proteins within ocular tissues. Our findings indicate that floRA is a reasonable surrogate for atRA binding to BSA, HDL, Apo A-I and RBP4 due to generally similar binding characteristics. Additionally, whole tissue tracer studies with floRA suggest it can be directly visualized in relevant ocular tissues, but further refinement is required.

### Binding affinities to putative carrier proteins

atRA and floRA exhibited binding affinity to BSA, HDL, Apo A-I, and RBP4, all proteins that could potentially serve as interstitial retinoic acid chaperones. The observed association constants were found to be very similar, in the range of 2-13 × 10^5^ M^-1^ (Table **1**; as determined by quadratic model fitting), with the exception of our positive control CRABP2, as expected and consistent with CRABP2’s role in retinoic acid binding and transport in the cytoplasm [34, 52]. In biological systems, when multiple proteins are present, the effective atRA transport and carrying capacity of a given protein depends on both the protein’s binding affinity for atRA and its local abundance, and it is therefore of interest to compare relative concentrations of putative carrier proteins.

- In humans, the serum concentration of albumin is typically 35-50 mg/ml, whereas in the interstitial space, it is 3-to 5-fold lower [53-55]. (We note that this is much larger than the serum concentration of atRA in humans, which typically ranges from 2.8 to 6.6 ng/ml [56, 57].) In mice, the serum albumin concentration is 10-30 mg/ml [58].
- Typical concentrations of Apo A-I in both human and mouse plasma are 1-2 mg/ml [59-62] and the average concentration in interstitial fluid is 8-to 12-fold lower [63].
- In humans, the concentration of retinol-RBP4 in plasma is maintained at 0.04–0.06 mg/ml, whereas in mice, it is around 0.02 mg/ml [64, 65]. It is also important to note that while the majority (∼90%) of RBP4 in the bloodstream exists as retinol-RBP4 complex, a small percentage is also estimated to circulate as free RBP4 (∼10%) [66-68].

In short, Apo A-I and RBP4 concentrations are one or more orders of magnitude lower than corresponding serum albumin concentrations. Considering the similar binding affinities of atRA to various protein binding partners, these data suggest that serum albumin would be expected to be the main binding partner for atRA and floRA transport in the serum [69] and in the extravascular space in the eye. At first glance this appears to differ somewhat from the conclusions of Summers, who identified Apo A-I as the dominant atRA-binding protein in the chick eye [19], consistent with earlier observations from Mertz and Wallman [15]. This may reflect a species difference (chicks vs. mammals), but it is also important to note that choroidal Apo A-I levels are increased at certain phases of myopigenic recovery and can also be upregulated by atRA [19]. Future studies of atRA signaling in myopigenesis should therefore consider both albumin and Apo A-I as atRA carriers, as well as assaying the local concentrations of each of these carriers at different phases of myopigenesis/recovery from a myopigenic stimulus.

### Comparison of protein binding of atRA vs. floRA

The binding characteristics of atRA and floRA were generally similar (e.g. binding affinities to BSA, HDL, Apo A-I, and RBP4) but also showed some differences. For example, we found that floRA bound to RBP1 whereas atRA did not (Figure **S5**). Although we observed floRA-RBP1 binding (K_a_ = 7.9 ± 0.4 × 10^5^ M^-1^, Table **1**), it was much weaker compared to the retinol-RBP1 and retinaldehyde-RBP1 binding in which K_a_ falls in the range of a few hundred × 10^5^ M^-1^ as reported in the literature [27, 28]. Further, we observed that floRA bound somewhat more weakly to RBP4 than did atRA (Figure **5**, Table **1**). Thus, although atRA and floRA share important similarities in their affinities for putative binding partners, they are not exact functional equivalents, i.e. floRA may not be a good atRA surrogate for transport studies in every circumstance. However, floRA appears to be a reasonable surrogate for determining the location of atRA-BSA complexes within tissue, and thus can be useful for mass transfer tracing studies of atRA in the eye.

### Initial studies using floRA as a tracer in ocular tissues

Based on the above, we used floRA-BSA conjugates as a surrogate for visualizing atRA transport in ocular tissues (Figure **7**). It was possible to observe floRA within ocular tissues but there were limitations with these experiments. These included a tendency for floRA-BSA to preferentially enter the tissue samples via the suprachoroidal space, possibly due to opening of this space during the tissue handling prior to incubation in the floRA-BSA solution. A second concern was the marked spatial heterogeneity observed in floRA signal within the tissue samples, perhaps due to segregation of floRA within scleral cells, particularly nuclei. Although this result is encouraging inasmuch it is consistent with the biology of atRA, which exerts its effects in the cell nucleus, it greatly complicates analysis of transport. Future analysis should consider subdividing tissue samples into cellular and acellular zones (based for example on vimentin or other secondary labeling of cells) and quantifying floRA signal in both cellular and acellular regions. A third complication was the pigmentation in the choroid, which possibly interfered with floRA signal and led us to focus preliminary quantitative analysis on the sclera. Despite these limitations which precluded rigorous quantitative analysis, our results clearly showed floRA-BSA penetration into both choroid and sclera, consistent with the idea that atRA is an element of the myopigenic retinoscleral signaling pathway.

### Limitations

This work is subject to several limitations. For example, the determination of association constants using fluorescence quenching makes some assumptions. In our analysis we mainly considered static quenching, where the protein and atRA (or floRA) form a nonfluorescent ground state complex. Quenching occurs due to energy transfer from the protein’s tryptophan residues to atRA upon complexation. The level of quenching in this scenario depends on the physical proximity of the binding domain and the tryptophan residue. It is possible that protein-ligand complexation takes place, but the binding domain is relatively far from the tryptophan residue(s), thereby impeding the energy transfer. In such a case, interpretation of the fluorescence quenching data as a measure of the actual binding affinity is limited. In addition, the fluorophore (in this case, proteins with intrinsic tryptophan fluorescence) can undergo dynamic quenching through random collision events with the same quencher responsible for static quenching. To overcome these limitations, additional studies using various spectroscopic methods (Fourier-transform infrared, Circular dichroism, fluorescence etc.) [22] alongside additional techniques such as microcalorimetry [70], and surface plasmon resonance [20, 71, 72] could be undertaken.

## Acknowledgements

We gratefully acknowledge Dr. Robert Hincapie for fruitful discussions during the initial experimental design. This work was supported by NIH EY033361 (SC, CRE, MP, MK), NIH F32 EY035573 and NIH T32 EY007092 (MBF), VA Rehab R&D RX003134 (MP) and the Georgia Research Alliance (CRE).

