## Supplemental Material for "Studies of all-*trans* retinoic acid transport in myopigenesis"

### Supporting Figures

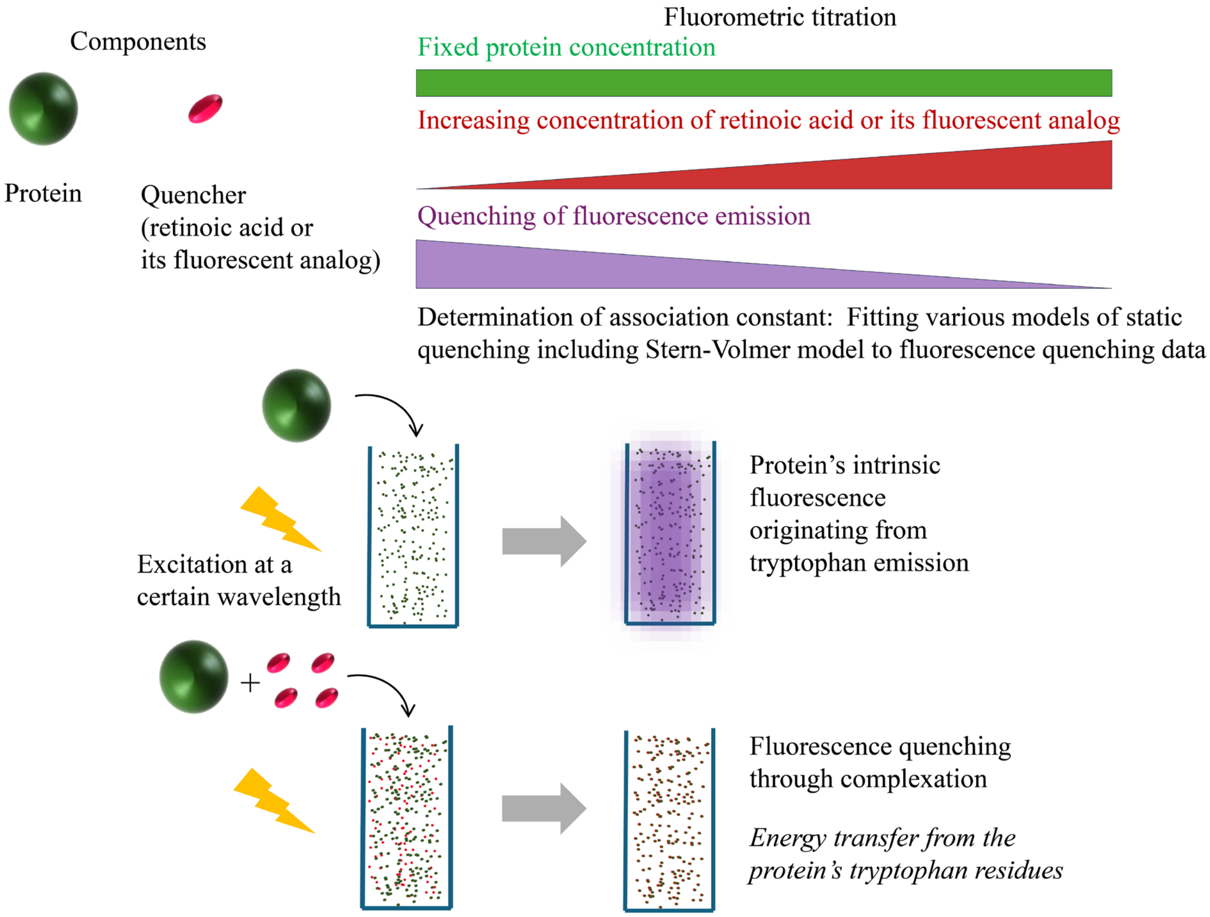


**Figure S1** Schematic diagram illustrating the experimental workflow for the fluorescence quenching experiments using carrier proteins and either atRA or its fluorescent analog, floRA.

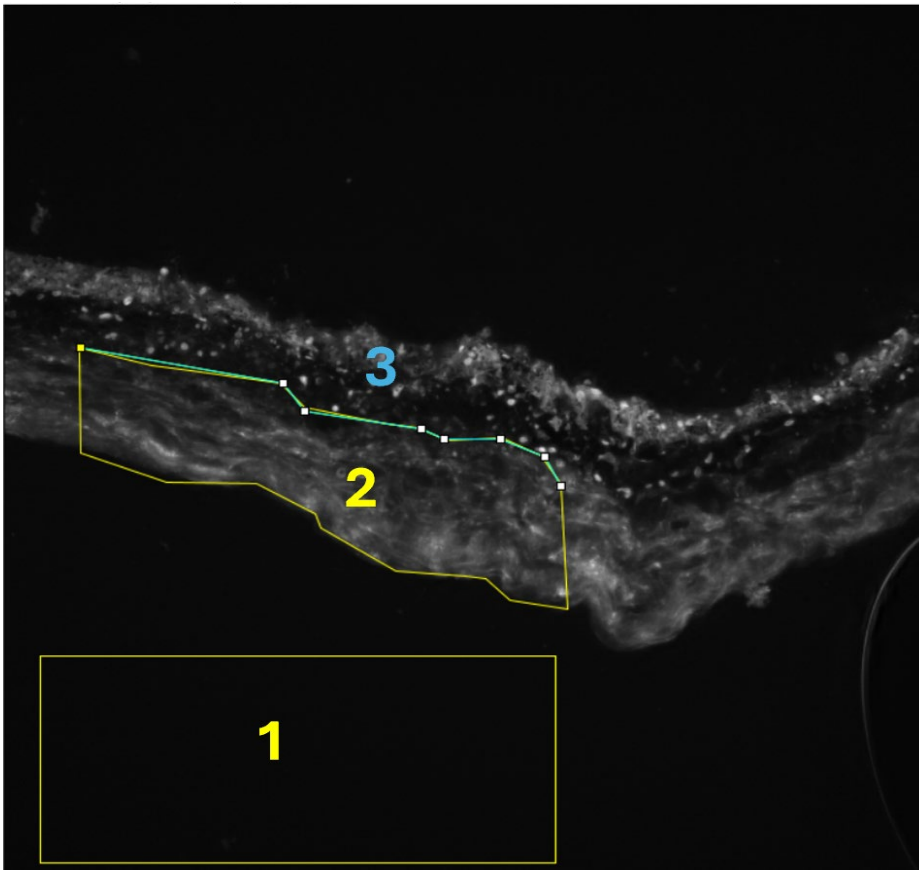


**Figure S2**: Florescent micrograph showing typical floRA labeling patterns and regions used for quantification of scleral fluorescence intensity. To quantify fluorescence and other metrics we use Fiji to carry out the following steps: (i) Define a rectangle separate from the sectioned tissue (region 1) and determine the mean fluorescence intensity in this rectangle, to be used as the background fluorescence level. (ii) Demarcate a scleral region (region 2) and determine the mean fluorescence intensity (scleral signal) and area of this region. (iii) Extract the length of curve 3, which forms the choroidal-scleral boundary of region 2. From these measurements we compute: (iv) the net fluorescence intensity in the sclera (average fluorescence intensity in region 2 – region 1), and (v) the “thickness” of the sclera, defined as the area of region 2 divided by the length of curve 3. The field of view in this image is 330.2 microns by 330.2 microns.

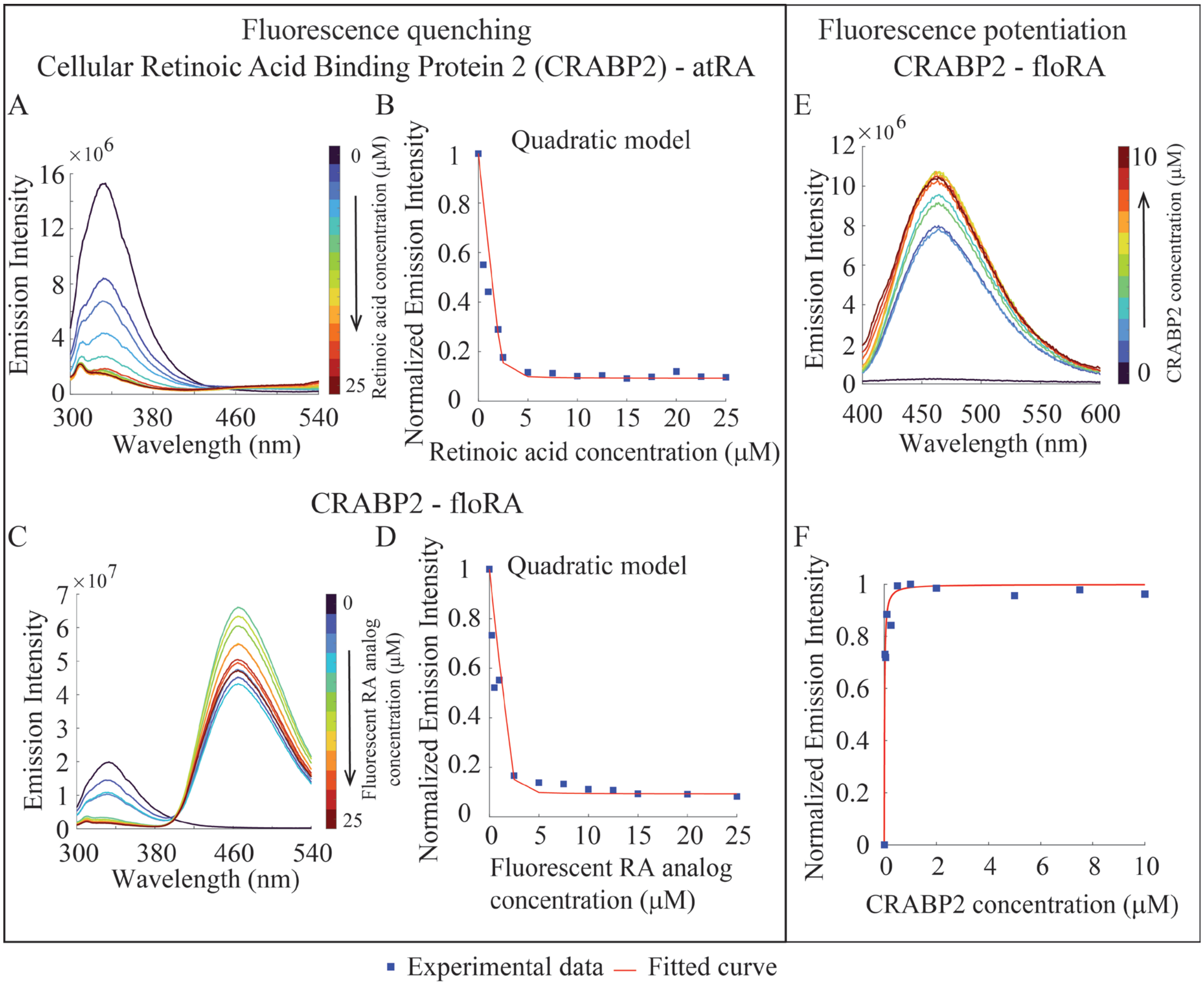


**Figure S3 Fluorometric investigations of CRABP2-retinoic acid binding. (A)** and **(C)** Fluorescence emission spectra of 2.4 μM CRABP2 in the presence of a range of concentrations (0-25 μM; colored curves) of atRA or floRA, respectively. The quenching of CRABP2 fluorescence with increasing concentrations of atRA or floRA is evident (see peak at 332 nm). There is an additional peak in the spectra in Figure **S3C** due to floRA having an absorbance peak around 280 nm which is absent for atRA (Figure **S4**). **(B)** and **(D)** Quadratic model fits to the peak emission intensities at 332 nm from the spectra in Figure **S3A** and **S3C,** respectively (see Methods). **(E)** Fluorescence emission spectra of 0.05 μM floRA in the presence of a range of concentrations of CRABP2 (0-10 μM; colored curves). With increasing concentration of CRABP2, the fluorescence intensity increases, reflecting the binding of more floRA to CRABP2. **(F)** Fit of the fluorescence emission peaks at 463 nm in Figure **S3E** to a standard Hill curve to estimate the association constant (see Methods). Panels **(A)-(F)** show typical results from 3 technical replicates.

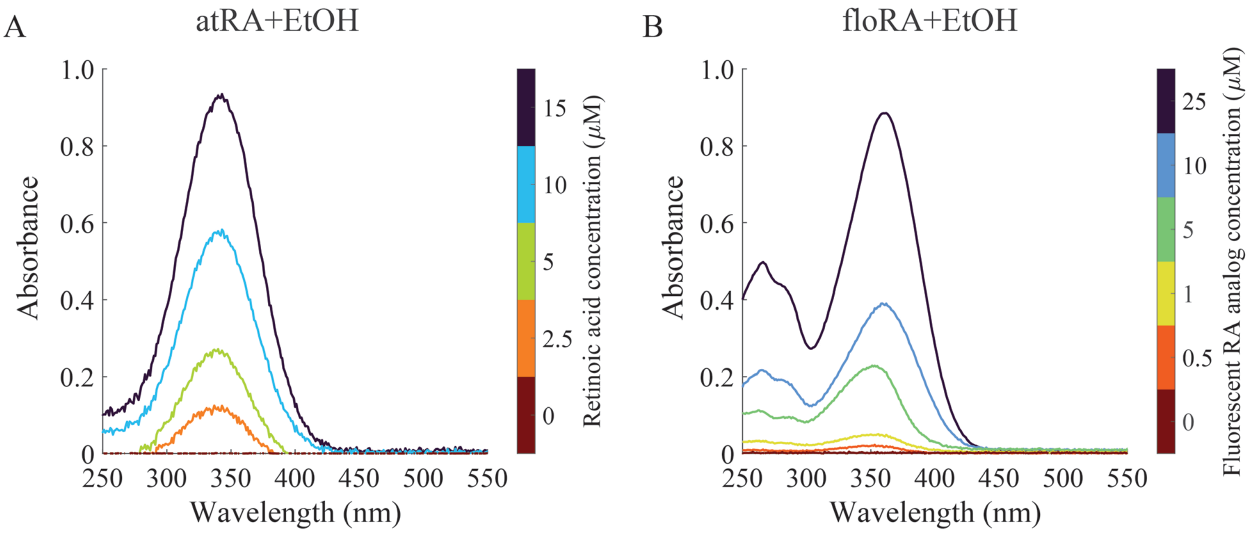


**Figure S4 (A)** UV-Vis absorbance spectra of 0-15 μM atRA in ethanol, showing an absorption peak around 340 nm **(B)** UV-Vis absorbance spectra of 0-25 μM floRA in ethanol showing absorption peaks around 265 nm, 280 nm, and 360 nm.

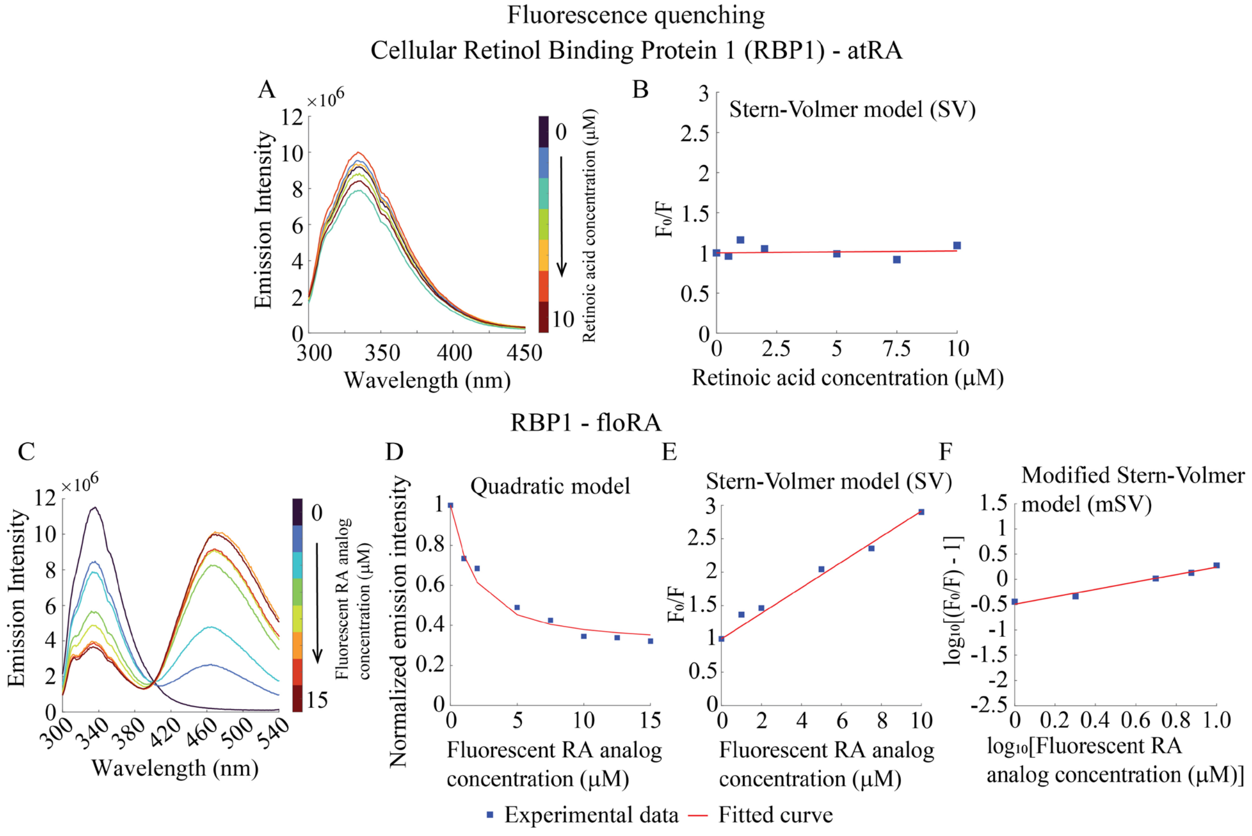


**Figure S5 Fluorometric investigations of RBP1-retinoic acid binding. (A)** and **(C)** Fluorescence emission spectra of 1.0 μM RBP1 in the presence of a range of concentrations (0-10 μM for atRA and 0-15 μM for floRA; colored curves) of atRA or floRA, respectively. The RBP1 fluorescence does not quench in presence of atRA, but quenches in presence of floRA (see peaks at 334 nm). There is an additional peak in the spectra in Figure **S5C** due to floRA having an absorbance peak around 280 nm which is absent for atRA (Figure **S4**). **(B)** and **(E)** SV model fits to the peak emission intensities at 334 nm from the spectra in Figure **S5A** and **S5C,** respectively. **(D)** Quadratic model fit to the peak emission intensities at 334 nm from the spectra in Figure **S5C**. **(F)** mSV model fit to the peak emission intensities at 334 nm from the spectra in Figure **S5C**. Panels **(A)-(F)** show typical results from 3 technical replicates.

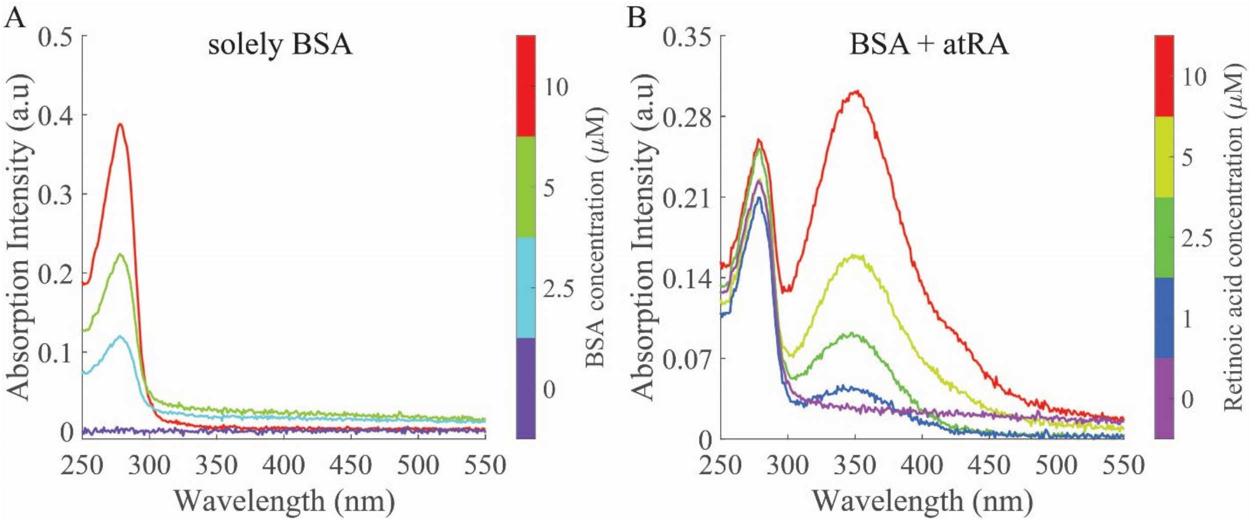


**Figure S6 (A)** UV-Vis absorbance spectra of 0-10 μM BSA in PBS. The absorbance peak appears around 280 nm **(B)** UV-Vis absorbance spectra of 5 μM BSA in the presence of 0-10 μM atRA in PBS (≤5% EtOH). Absorbance peaks were observed around 280 nm (for BSA) and 350 nm (for atRA).

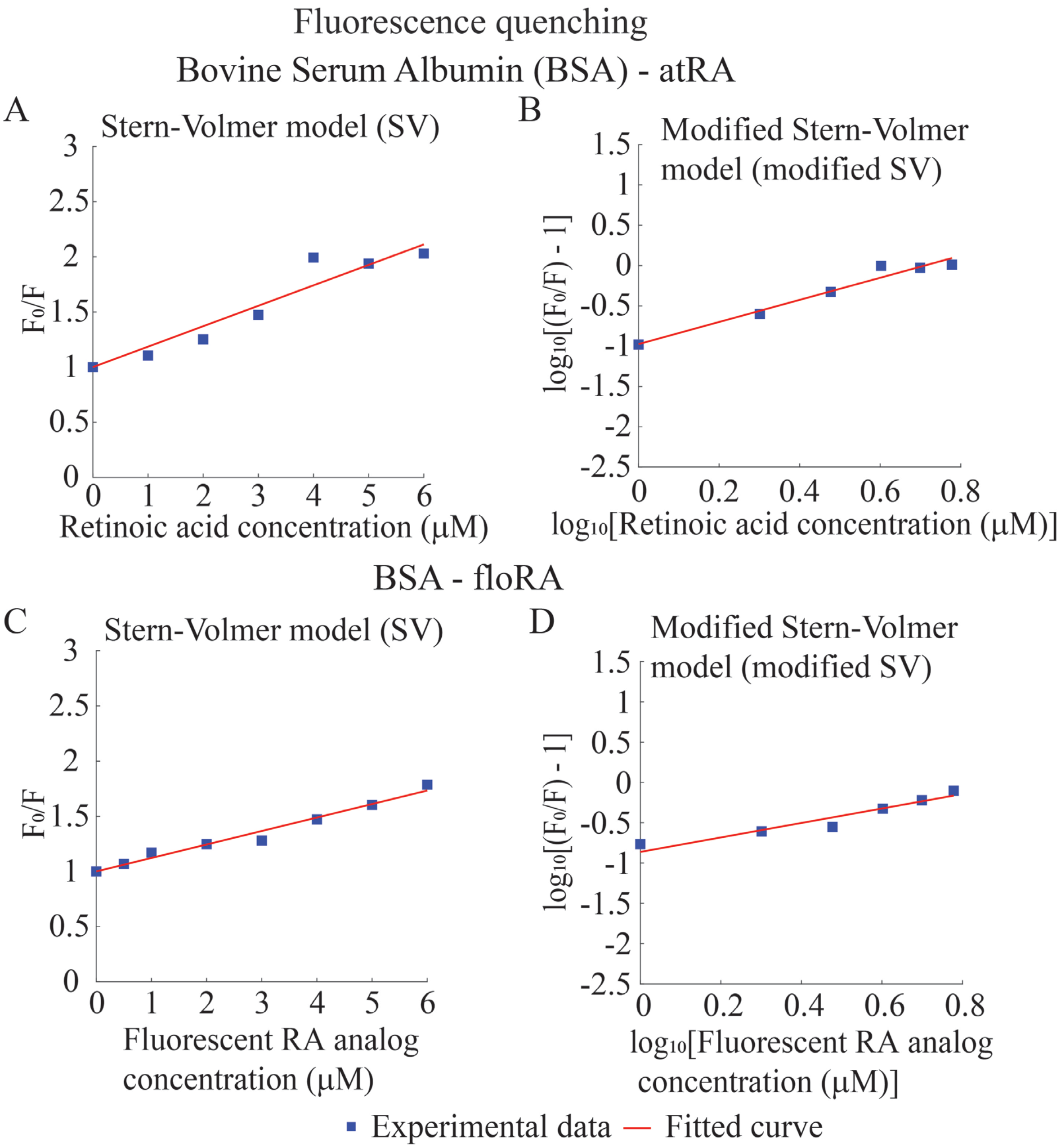


**Figure S7 Estimation of association constants for BSA-atRA/floRA complexation (A-B)** Stern-Volmer (SV) **(A)** and modified SV **(B)** model fits to the peak emission intensities at 336 nm from the spectra in Figure **2A. (C-D)** SV **(C)** and modified SV **(D)** model fits to the peak emission intensities at 336 nm from the spectra in Figure **2C.**

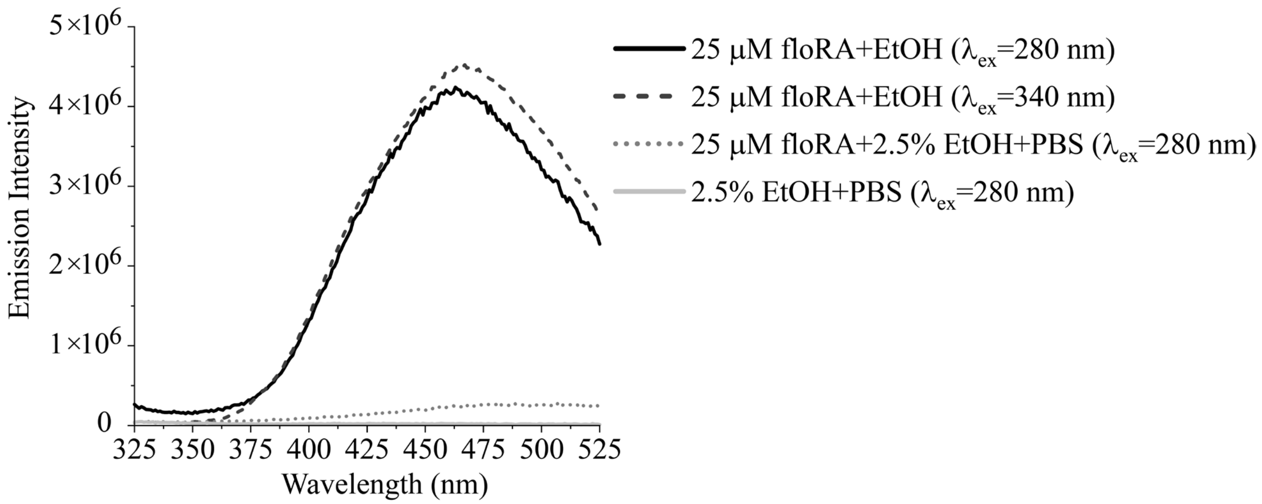


**Figure S8 Comparison of the fluorescence spectra of floRA in aqueous vs. non-aqueous solvents**. 25 µM floRA shows fluorescence emission in the organic solvent EtOH, but not in aqueous environment (e.g. PBS).

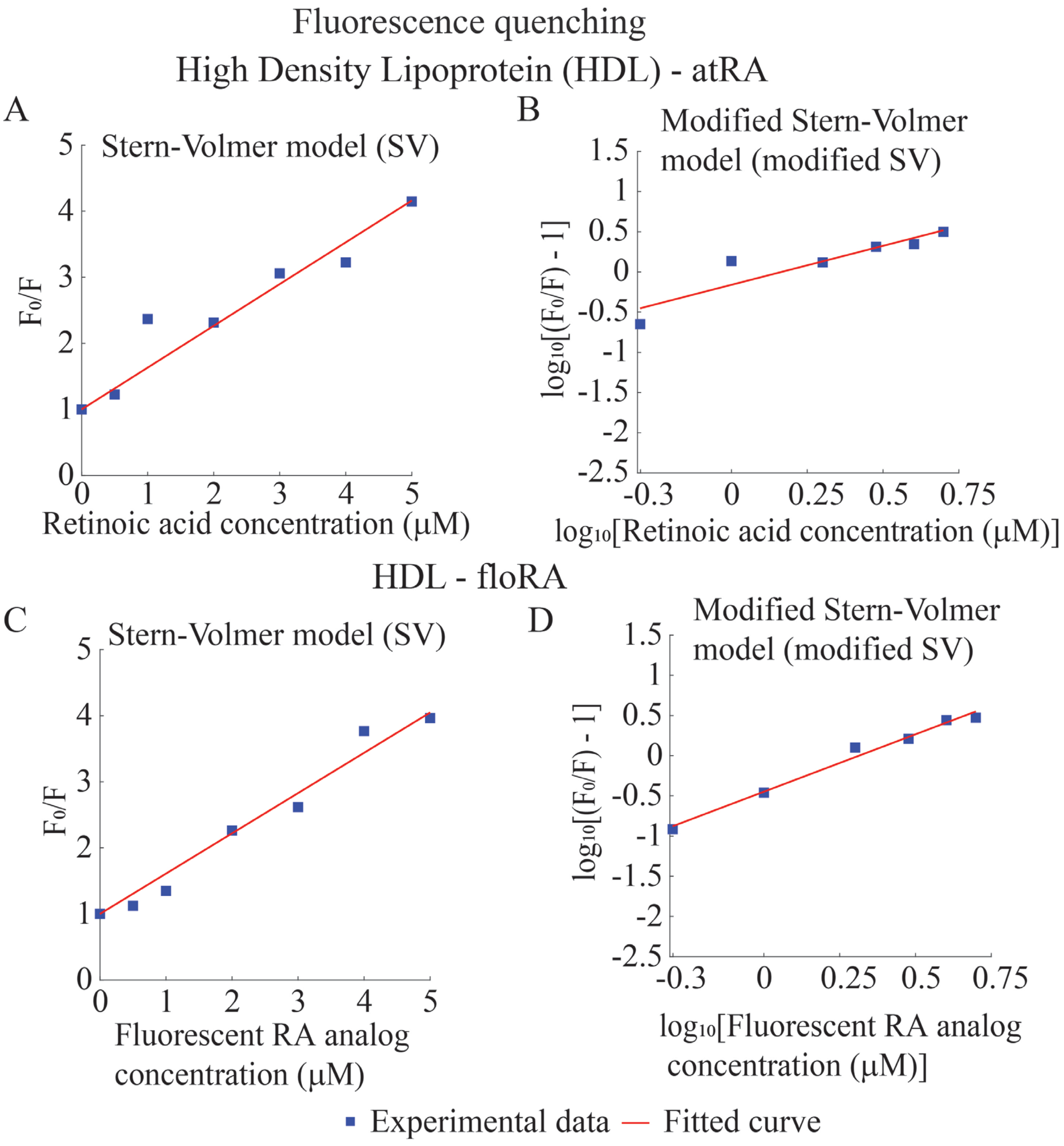


**Figure S9 Estimation of association constants for HDL-atRA/floRA complexation (A-B)** Stern-Volmer (SV) **(A)** and modified SV **(B)** model fits to the peak emission intensities at 321 nm from the spectra in Figure **3A. (C-D)** SV **(C)** and modified SV **(D)** model fits to the peak emission intensities at 321 nm from the spectra in Figure **3C.** Panels **(A)-(D)** show typical results from 3 technical replicates.

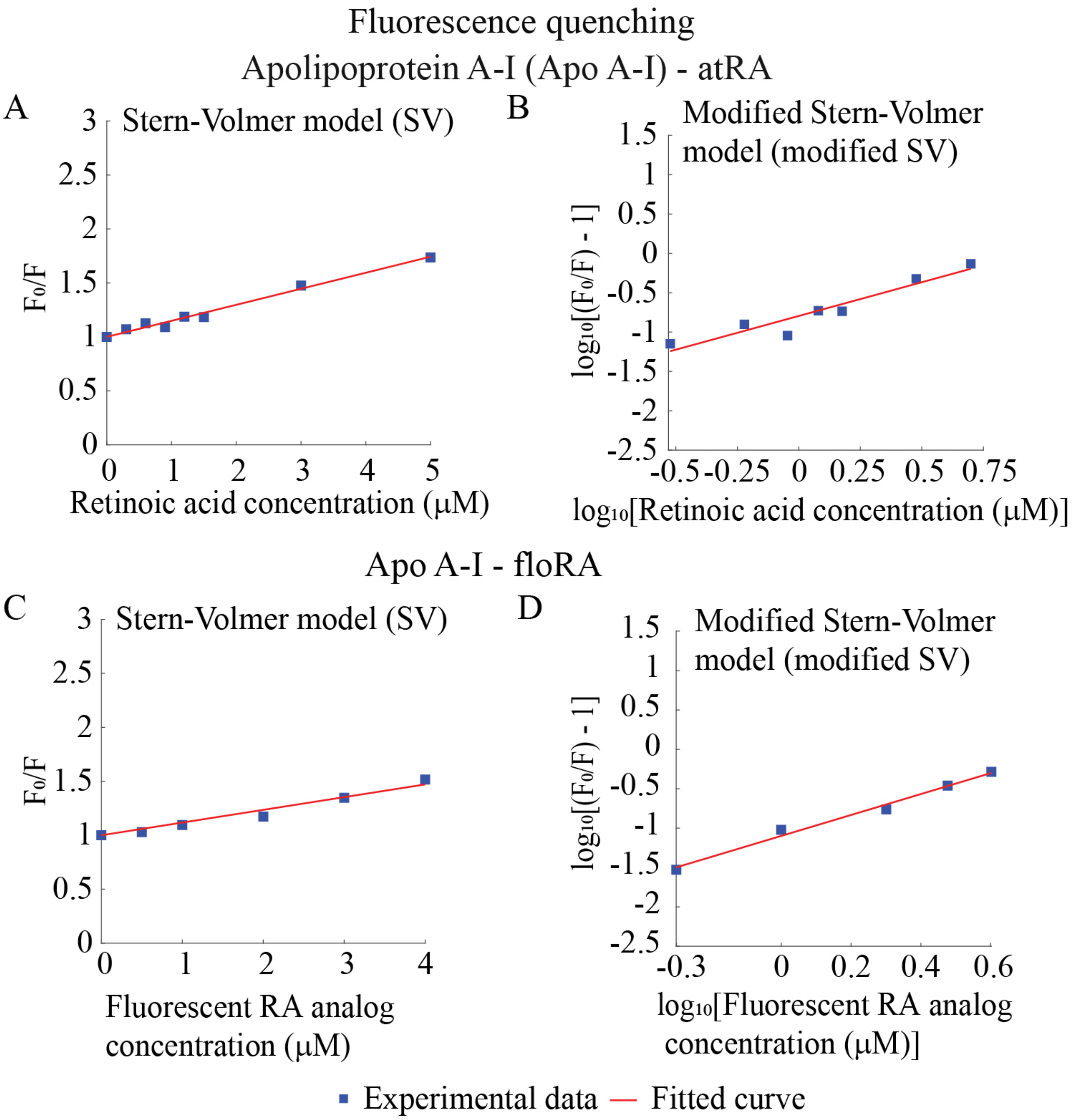


**Figure S10 Estimation of association constants for Apo A-I-atRA/floRA complexation (A-B)** Stern-Volmer (SV) **(A)** and modified SV **(B)** model fits to the peak emission intensities at 332 nm from the spectra in Figure **4A. (C-D)** SV **(C)** and modified SV **(D)** model fits to the peak emission intensities at 332 nm from the spectra in Figure **4C**. Panels **(A)-(D)** show typical results from 3 technical replicates.

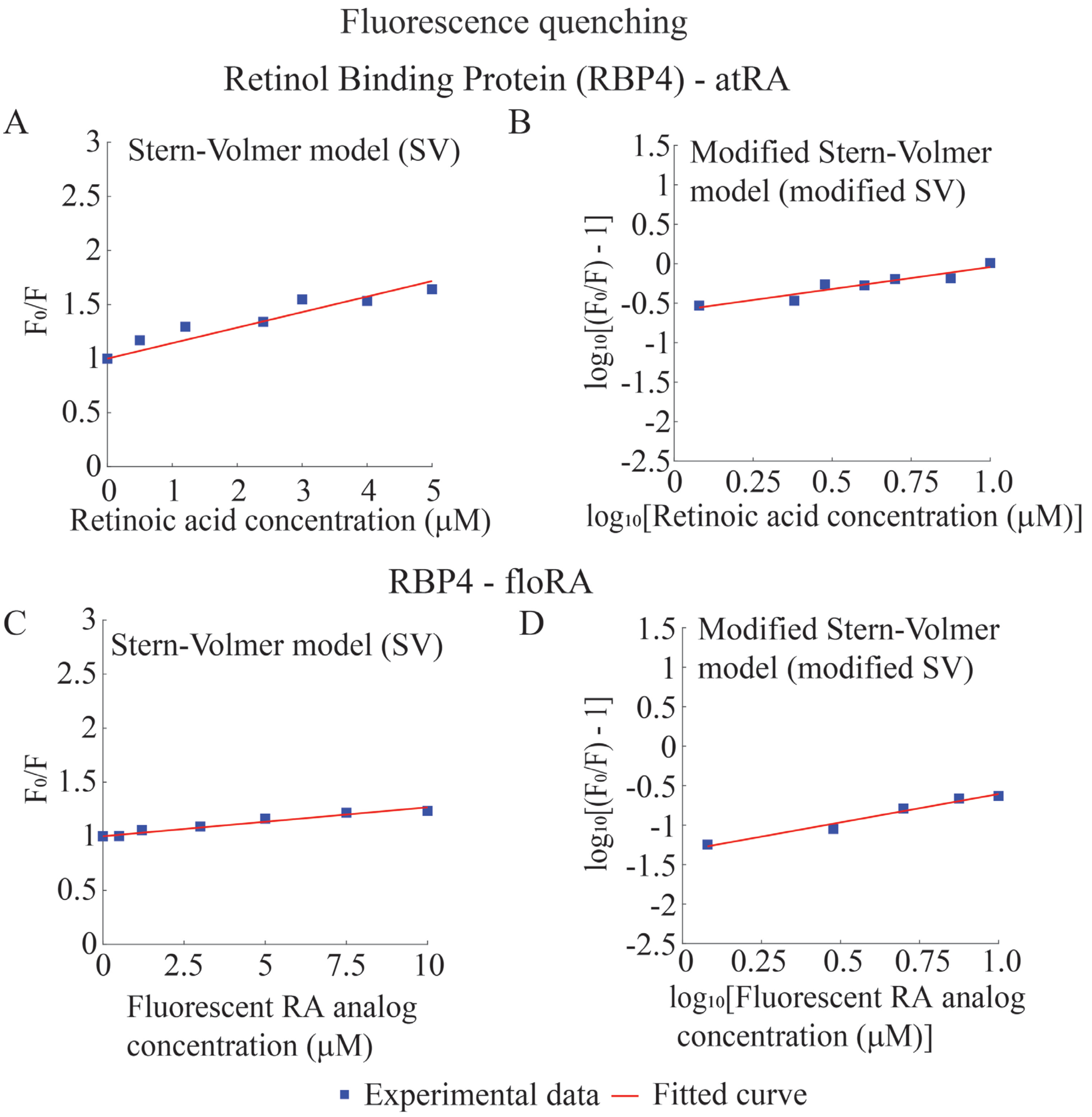


**Figure S11 Estimation of association constants for RBP4-atRA/floRA complexation (A-B)** Stern-Volmer (SV) **(A)** and modified SV **(B)** model fits to the peak emission intensities at 334 nm from the spectra in Figure **5A. (C-D)** SV **(C)** and modified SV **(D)** model fits to the peak emission intensities at 334 nm from the spectra in Figure **5C.** Panels **(A)-(D)** show typical results from 3 technical replicates.

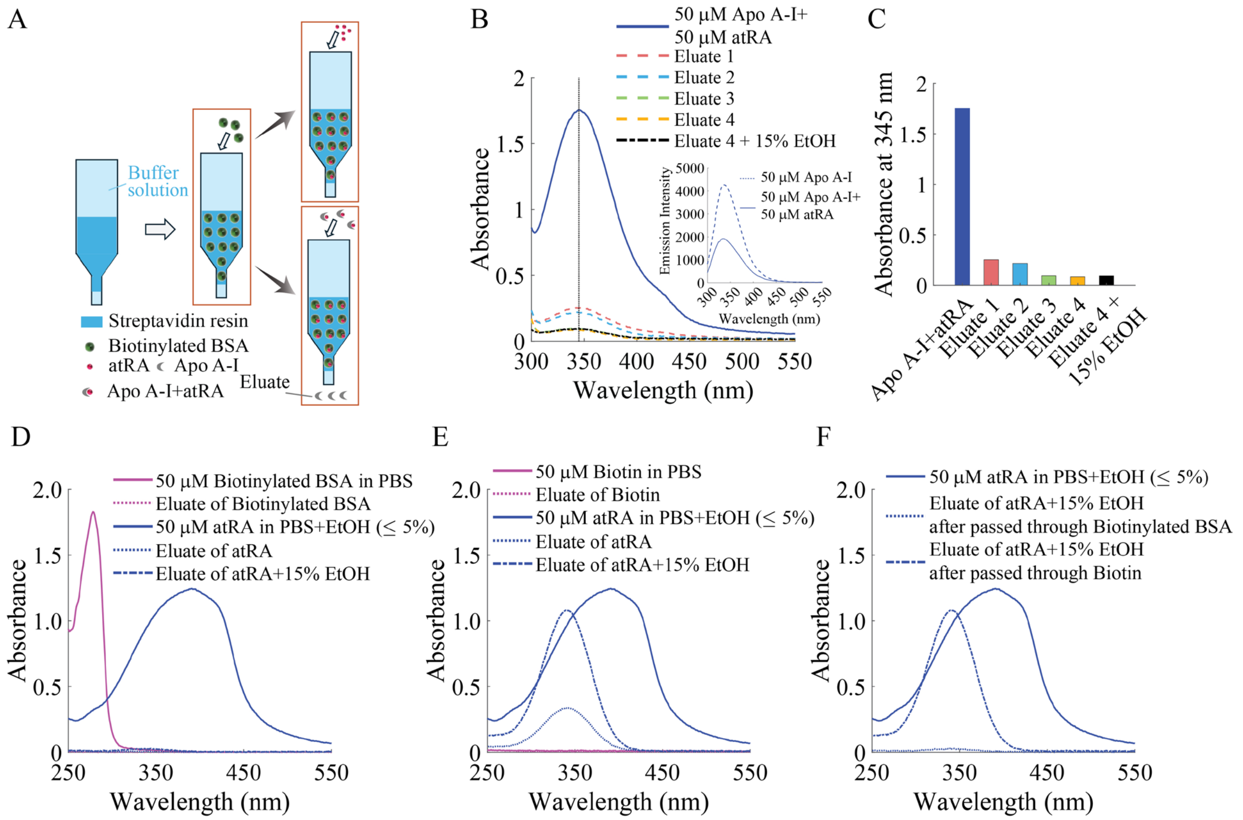


**Figure S12 Additional experiments with atRA in presence of biotin, biotinylated BSA and Apo A-I (A)** Schematic illustration of the experimental design. Solely atRA and Apo A-I-atRA mixture were separately passed through the biotinylated BSA coated streptavidin resin packed columns. The corresponding eluates were collected for analysis. **(B)** UV-Vis absorbance spectra of Apo A-I-atRA mixture and subsequent eluates. (Inset) A reference emission spectra of 50 µM Apo A-I solution in presence and absence of 50 µM atRA (excitation wavelength 280 nm). (**C**) Absorbance maxima of Apo A-I-atRA mixture and subsequent eluates at 345 nm (as depicted in Figure **S12B**). **(D-F)** UV-Vis absorbance spectra acquired from the control experiments with atRA passed through Biotin/Biotinylated BSA bound streptavidin resin and the corresponding eluates. Some of the dotted/dashed curves are difficult to visualize as they partially overlap the horizontal axis.

### Supporting Tables

| **Protein** | **Commercial source** | **Lot no.<br>Catalog no.** | **Received as** |
|---|---|---|---|
| High Density Lipoproteins (HDL), Human Plasma | Athens Research and Technology, GA, USA | HDL2023-09<br>12-16-080412 | Liquid; 5 mg in 403 µL of 150 mM NaCl (pH 7.4) 0.01% EDTA |
| | | HDL2023-09<br>12-16-080412 | Liquid; 5 mg in 455 µL of 150 mM NaCl (pH 7.4) 0.01% EDTA |
| Apolipoprotein AI (Apo A-I), Human Plasma | Athens Research and Technology, GA, USA | APOA12022-01<br>16-16-120101 | Frozen liquid; 1 mg in 625 µL 10 mM $NH_4HCO_3$ (pH 7.4) |
| Retinol Binding Protein (RBP4), Human Plasma | Athens Research and Technology, GA, USA | RBP2020-01<br>16-16-180216 | Lyophilized solid.<br>Pre-lyophilized: 100 µg in 39.1 µL 50 mM Tris with 100 mM NaCl (pH 7.4) |
| Bovine Serum Albumin (BSA) | Fisher Scientific, USA | 110466<br>BP1605-100 | Refrigerated, lyophilized solid |

**Table S1** Sources of purchased proteins, and their concentrations as received.

| Protein | Ligand | Protein concentration (µM) | Ligand concentration (µM) | Excitation wavelength (nm) | Emission peak (nm) monitored at |
|---|---|---|---|---|---|
| BSA | atRA | 6 | 0-25 | 280 | 336 |
| | floRA | 6 | 0-25 | 280 | 336 |
| HDL | atRA | 0.028 mg/ml | 0-25 | 280 | 321 |
| | floRA | 0.028 mg/ml | 0-25 | 280 | 321 |
| Apo A-I | atRA | 1.2 | 0-25 | 280 | 332 |
| | floRA | 1.2 | 0-25 | 280 | 332 |
| RBP4 | atRA | 1.2 | 0-15 | 280 | 334 |
| | floRA | 1.2 | 0-15 | 280 | 334 |
| CRABP2 | atRA | 2.4 | 0-25 | 280 | 332 |
| | floRA | 2.4 | 0-25 | 280 | 332 |
| RBP1 | atRA | 1.0 | 0-10 | 280 | 334 |
| | floRA | 1.0 | 0-15 | 280 | 334 |

**Table S2** Experimental details for protein-atRA and protein-floRA fluorescence quenching experiments.

| Protein | Ligand | Protein concentration (μM) | Ligand concentration (μM) | Excitation wavelength (nm) | Emission peak monitored (nm) |
|---|---|---|---|---|---|
| BSA | floRA | 0-25 | 0.05 | 340 | 461 |
| HDL | floRA | 0-0.7 mg/ml | 0.05 | 340 | 437 |
| Apo A-I | floRA | 0-35 | 0.05 | 340 | 467 |
| RBP4 | floRA | 0-18 | 0.025 | 340 | 466 |
| CRABP2 | floRA | 0-10 | 0.05 | 340 | 463 |

**Table S3** Experimental details for protein-floRA fluorescence potentiation experiments.

| Fluorescence Potentiation | | |
|---|---|---|
| Protein | Ligand | Hill curve fit |
| | | $K_a$ (× $10^5$ $M^{-1}$) |
| BSA | floRA | 11.8 ± 1.2 |
| HDL | floRA | 10.3 ± 3.5 |
| Apo A-I | floRA | 2.5 ± 0.1 |
| RBP4 | floRA | 2.2 ± 0.5 |
| CRABP2 | floRA | 539.3 ± 169.6 |

**Table S4** Association constants (mean ± SD) estimated from fluorescence potentiation experiments with floRA and several proteins.

**Additional details on the determination of protein-atRA and protein-floRA association constants and stoichiometry from fluorescence quenching data**

*Stern-Volmer model:* The Stern-Volmer model has been widely utilized to evaluate the stoichiometry and binding constants associated with static quenching, where the fluorescently active molecule forms a complex with a quencher molecule [1-4]. In the context of retinoic acid-protein complexation, the protein molecule having tryptophan fluorescence plays the role of the fluorescently active molecule (P) whereas atRA (or floRA) serves as the quencher (Q). This model assumes an equilibrium reaction P + Q ⇌ PQ, with PQ being the nonfluorescent ground state complex. The fluorescence intensity is then expressed in relation to the concentration of the quencher by the classic Stern-Volmer equation [5]:

$F_0/F = K_a [Q] + 1$ (1)

Here F is the fluorescence intensity in the presence of Q, $F_0$ is the fluorescence intensity in the absence of Q, $K_a$ is the Stern-Volmer constant, and [Q] is the quencher molar concentration, i.e. either atRA or floRA in this study. This model assumes that the quencher forms a complex with the protein and the quenching constant $K_a$ can be considered equivalent to the binding constant of the complexation reaction, as static quenching results from the formation of a non-emissive dark complex between the fluorescently active molecule P and the quencher Q. A modified version of the Stern-Volmer model considers an equilibrium reaction, $P + nQ \rightleftharpoons PQ_n$, where n units of quencher Q form a complex with a single fluorescently active molecule P. In this scenario, the modified Stern-Volmer equation is:

$$\log_{10}(F_0/F - 1) = \log_{10} K_a + n\log_{10} [Q] \qquad (2)$$

The 'fit' function in MATLAB was used to fit the fluorescence quenching data according to Equation (1) or (2).

*Quadratic model:* This model considers the same equilibrium reaction, $P + Q \rightleftharpoons PQ$. This model of static quenching, in which quenching occurs through formation of a complex between the fluorescently active molecule and the quencher, assumes 1:1 binding stoichiometry between the protein and the quencher molecules. It also assumes that the quencher (atRA or floRA) is either free in solution or bound to the protein and the fluorescence of the free quencher is zero [6-8]. Considering that the fluorescence quenching is proportional to the degree of binding (ratio of protein-quencher complex and total protein concentration $[P_T]$), the fluorescence emission intensity in a protein-quencher solution can be expressed as:

$$F = F_0 - \frac{1+([P_T]+[Q])K_a - \sqrt{([P_T]-[Q])^2 K_a^2 + 2([P_T]+[Q])K_a + 1}}{2[P_T]\,K_a}(F_0 - F_\infty) \qquad (3)$$

Here, $[P_T]$ is the total protein concentration; [Q] denotes the total quencher concentration (atRA or floRA), $K_a$ denotes the association constant and $F_\infty$ represents the fluorescence signal after the protein has been completely saturated with ligand. This model can deal with nonlinearity in fluorescence quenching, especially at high quencher concentrations, whereas the traditional Stern-Volmer model cannot. To fit Equation (3) to the fluorescence data, we performed a nonlinear least squares curve fitting using MATLAB's 'lsqcurvefit' function.

### Additional notes on the analysis of HDL-atRA and HDL-floRA fluorometric titration experiments

The HDL solution we used (Table **S1**) was a mixture of cholesterol, triglyceride and multiple proteins including Apo A-I (molecular mass 28 KDa) and Apo A-II (molecular mass 17.4 kDa), composed of 45-55% lipid with the remainder being proteins. Since this solution was a multi-protein mixture with an unknown fraction of each protein, we report its effective total protein concentration as a mass, rather than a molar concentration. In HDL-atRA quenching experiments (Figure **3** and Figure **S9**), we kept the HDL concentration fixed at 0.028 mg/ml and varied the atRA concentration over the range 0 – 25 µM (Figure **3A-3B**, Table **S2**). However, to fit the quadratic model, we required the molar protein concentration as an input parameter (see the previous section), which was unknown. We therefore treated the effective protein concentration as a free parameter in the model fitting [6]. Using this approach, from HDL-atRA quenching experiments, we obtained an effective protein concentration of 0.73 ± 0.06 µM corresponding to the HDL concentration of 0.028 mg/ml. Of note, a concentration of 0.028 mg/ml Apo A-I (molecular weight 28 kDa) corresponds to approximately 1 µM, aligning closely with the estimated effective protein concentration within HDL. The same approach yielded an effective protein concentration of approximately 2 μM corresponding to same HDL concentration from the HDL-floRA quenching experiments (Figure **3C-3D**, Table **S2**). In the fluorescence potentiation experiments, the floRA concentration was fixed at 0.05 μM and we varied the HDL concentration over the range 0 – 0.705 mg/ml (Figure **3E-3F**, Table **S3**). Subsequently, we fitted a standard Hill curve to the peak emission intensities monitored at 437 nm (Figure **3F**). We then used the effective protein concentration of 0.028 mg/ml within the HDL solution, which is equivalent to approximately 2 μM (derived from quadratic model fits to HDL-floRA quenching experiments), to estimate the HDL-floRA association constant as $10.3 \pm 3.5 \times 10^5\ M^{-1}$ (Table **S4**).
